# Dim Green Light Enables Day-and-Night Monitoring of Leaf Movements

**DOI:** 10.64898/2026.05.08.723725

**Authors:** Eva Herrero, Alison R. Gill, Samalka Wijeweera, Daniel N. Ginzburg, John D. Stamford, Andreas Antoniades, Jennifer R. Bromley, Jenny C. Mortimer, Matthew Gilliham, A. Harvey Millar, Alex A.R. Webb

## Abstract

Understanding plant growth dynamics requires imaging across day-and-night cycles to quantify growth, movement and development in the aerial plant body and to capture the rhythmic nature of these processes. This requires imaging in light during the day and in darkness at night without perturbing plant physiology. Nighttime imaging has typically depended on infrared (IR) illumination, producing monochrome datasets that require specialised hardware and separate analysis pipelines when combined with daytime RGB imaging. Here, we evaluated very low-intensity green (dimG) illumination from standard LEDs as a practical alternative for colour-consistent nighttime imaging and assessed its physiological impact in *Arabidopsis thaliana* and *Lactuca sativa* (lettuce). We show that high resolution colour images can be obtained under dimG using low- cost cameras, with sufficient consistency between full-spectrum and dimG images to allow direct comparison and unified image analysis. We show that very low-fluence green light (<0.5 μmol m^-2^ s^-1^) does not sustain circadian oscillations of gene activity under continuous exposure and does not perturb rhythms when applied during the dark phase of diel cycles. DimG imaging enabled accurate detection of diel leaf movement profiles in *Arabidopsis* circadian mutants, revealing genotype-specific phase differences under varying photoperiods. In lettuce, dimG pulses and continuous dimG enabled accurate quantification of diel leaf movement without affecting growth, stomatal opening, electron transport rate or chlorophyll content. Motion profiles under continuous dimG mirrored those under darkness. Our findings establish dim green illumination as a cost-effective solution for night-time imaging, simplifying phenotyping workflows with minimal impact on physiology.

## Introduction

Continuous imaging across day-night (diel) cycles enables high-resolution quantification of growth dynamics, circadian leaf movements, developmental transitions, and stress-related physiological responses (Dornbusch et al., 2014; Olas et al., 2020). Automated phenotyping systems, typically equipped with top- or side-mounted RGB cameras in controlled-environment chambers, acquire images at fixed intervals throughout both light and dark phases. However, achieving this continuity is technically challenging because RGB sensors typically require visible light illumination, which can activate plant photoreceptors and alter physiological processes during the dark phase (Huq et al., 2024). To circumvent this, infrared (IR) illumination using wavelengths which lie beyond the sensitivity range of the major plant photoreceptors (Battle & Jones, 2020) is often used and detection is performed with IR-sensitive cameras. While effective, this approach introduces complexity: standard LED panels do not emit IR, necessitating dedicated IR light sources and cameras (Bours et al., 2012; Olas et al., 2020). Moreover, daytime RGB images derive their pixel values from three visible-light colour channels shaped by pigment absorption, whereas nighttime IR images are effectively monochromatic and dominated by leaf structural scattering (Peñuelas & Filella, 1998; Ustin & Jacquemoud, 2020). Therefore, switching between these modalities produces abrupt, non-biological discontinuities in leaf brightness and contrast that interfere with continuous movement quantification. These limitations highlight the need for alternative strategies that maintain colour fidelity and avoid complications of combining RGB and monochromatic datasets for image analysis, while minimizing physiological disruption.

Low-intensity green illumination is generally regarded as minimally disruptive for nighttime sampling of leaf tissue (Johansson et al., 2014; Sorkin & Nusinow, 2022). Based on this premise, green wavelengths emitted by standard LEDs could provide nocturnal lighting at low intensity while enabling visible light image acquisition comparable to daytime conditions. However, green light is rarely employed for nighttime plant imaging (Zhang et al., 2012), likely due to concerns about physiological effects from continuous exposure. Although green light typically triggers weaker photoreceptor-mediated responses than red, far-red, or blue light, even low-fluence green illumination (7 µmol m⁻² s⁻¹) has been reported to influence circadian gene expression rhythms in *Arabidopsis* (Battle & Jones, 2020). If very low green-light exposure (<0.5 µmol m⁻² s⁻¹) does not meaningfully alter physiology while still supporting robust visible light imaging, it could make green light a cost-effective, low-processing solution for nocturnal imaging in agricultural and laboratory settings.

Leaf movement provides a sensitive and non-invasive proxy for evaluating the physiological impact of nocturnal illumination. Many plant species exhibit rhythmic changes in leaf angle and petiole position driven by the circadian clock, which highly responsive to light quality and intensity (Dornbusch et al., 2014; Oskam et al., 2024). These nyctinastic movements have long served as indicators of circadian integrity, as alterations in amplitude, phase, or period often reflect light-induced perturbations of clock function. Leaf movement has been widely used to investigate circadian rhythms under continuous light, a condition in which leaf movements are driven solely by the internal circadian clock rather than external cues (Edwards & Millar, 2007; Lou et al., 2022). Fewer studies have quantified leaf-movement dynamics under diel light cycles, including laser-scanning analysis of single leaves (Dornbusch et al., 2014) and time-lapse imaging with infrared cameras with semi-automated analysis of single leaves (Bours et al., 2012; Oskam et al., 2024) or whole-plants (Apelt et al., 2015). Together, these studies demonstrate the feasibility of high-resolution monitoring of leaf movements under low-light conditions. However, reliance on expensive instrumentation, such as laser scanning and light-field cameras, or analysis restricted to individual leaves highlights the need for a more accessible approach that is suited to whole-plant or multi-plant array imaging.

Here, we introduce a green light-based imaging strategy that overcomes the limitations of IR-based approaches by enabling colour-consistent visible light imaging throughout the diel cycle. We demonstrate that very low-fluence green illumination (<0.5 µmol m⁻² s⁻¹) provides sufficient light for high-quality image acquisition while minimising effects on physiology, as evidenced by unaltered circadian-regulated processes and growth patterns. Using *Arabidopsis* and lettuce as model systems, we further show that this approach supports precise quantification of diel leaf movement rhythms, establishing very low-intensity green illumination (dimG) imaging as a practical and minimally disruptive solution for continuous phenotyping.

## Results

### Dim green light does not sustain circadian oscillations of gene activity or photosynthetic activity

To evaluate whether dimG is an appropriate illumination source for nighttime imaging, we tested if the lowest intensity of green light emitted by our LED panel (dimG, 0.3 µmol m⁻² s⁻¹) affects light signaling. Promoter:luciferase (LUC) reporters are widely used to monitor circadian gene activity in plants, providing a non-invasive, high-resolution readout of transcriptional rhythms *in vivo* (Harmer, 2025). These reporters have robust oscillations under continuous light, where photoreceptor signalling sustains rhythm amplitude, defined as the difference between the peak and the mid-point of the oscillation. In continuous darkness, however, circadian rhythms typically show progressive damping due to the absence of light-dependent inputs that maintain amplitude (Haydon et al., 2017; Jones et al., 2015). A commonly used reporter, *pCCA1::LUC*, in which the promoter of the circadian oscillator component *CCA1 (CIRCADIAN CLOCK ASSOCIATED 1)* drives LUCIFERASE expression, is a well-established proxy for circadian clock activity (Airoldi et al., 2019). Previously, it was reported that green light at fluences >7 µmol m⁻² s⁻¹ enhance circadian oscillations of *CCA1* promoter activity in *Arabidopsis* (Battle & Jones, 2020). We therefore examined whether 0.3 µmol m⁻² s⁻¹ elicits a similar response.

We measured circadian oscillations of bioluminescence in *Arabidopsis thaliana* seedlings carrying *pCCA1::LUC*. Under diel cycles, nighttime illumination with either 10 or 0.3 µmol m⁻² s⁻¹ green light (Fig. 1A) did not alter *pCCA1::LUC* oscillations compared with darkness (Fig. 1B, D and F, 0 - 48 hours). Under continuous conditions, we assessed rhythm robustness using the Relative Amplitude Error (RAE), which estimates how well a fitted co-sine curve matches the measured signal. RAE values below 0.5 indicate a robust circadian rhythm (Kolmos et al., 2011). Under continuous darkness (DD), the *pCCA1::LUC* rhythms damped rapidly and became arrhythmic (Fig. 1B-C). Continuous exposure to 10 µmol m⁻² s⁻¹ green light partially rescued rhythms, with 42% of seedlings maintaining oscillations with a RAE < 0.5 (Fig. 1D-E), consistent with a previous study (Battle & Jones, 2020). On the other hand, continuous exposure to 0.3 µmol m⁻² s⁻¹ green light failed to sustain circadian oscillations, and rhythms damped similarly to DD (Fig. 1F-G). Together, these results demonstrate that very low-fluence green light (dimG) does not influence *CCA1* promoter-driven rhythms under either diel or continuous conditions, supporting its use as a physiologically neutral nighttime illumination source for imaging of diel and circadian rhythms.

**Figure 1.**
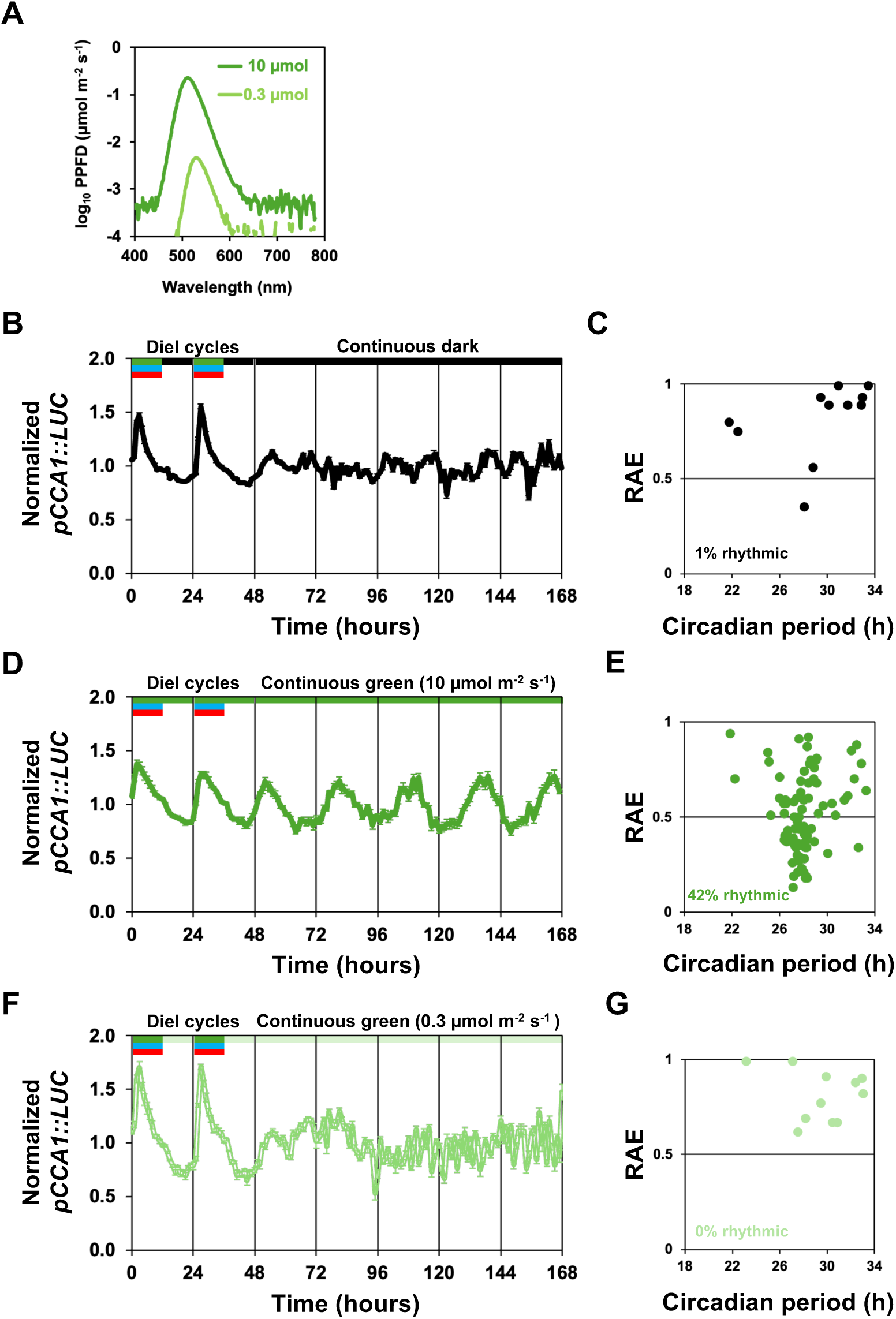
Dim green light does not sustain circadian oscillations of promoter activity in *Arabidopsis*. **(A)** Emission spectra of green light LEDs for 10 µmol m⁻² s⁻¹ or 0.3 µmol m⁻² s^⁻1^. Average *pCCA1::LUC* normalized bioluminescence. **(B)** Diel cycles with dark night followed by continuous dark (DD). **(C)** RAE versus circadian period length of curves from 72 to 168 h under DD. **(D)** Diel cycles with 10 µmol m⁻² s⁻¹ green light at night followed by continuous 10 µmol m⁻² s⁻¹ green light (average of rhythmic plants only). **(E)** RAE versus circadian period length of curves from 72 to 168 h under continuous 10 µmol m⁻² s⁻¹ green light. **(F)** Diel cycles with 0.3 µmol m⁻² s⁻¹ green light at night followed by continuous 0.3 µmol m⁻² s⁻¹ green light. **(G)** RAE versus circadian period length of curves from 72 to 168 h under continuous 0.3 µmol m⁻² s⁻¹ green light. Seedlings were entrained for 10 days (12h day/ 12 dark) with 70 µmol m⁻² s⁻¹ of white light. Plants with RAE < 0.5 were considered rhythmic. In B, D, F colour lines at the top of the graphs represent periods of darkness (black), 10 µmol m⁻² s⁻¹ green light only (dark green), 0.3 µmol m⁻² s⁻¹ (light green) and full RGB light (red, green and blue).

To further assess the physiological impact of dimG in Arabidopsis, we measured photosynthetic activity (Fig. S1) following one hour of exposure to complete darkness, green light at 0.3 or 10 µmol m⁻² s⁻¹, and full-spectrum RGB light (Figs. 1A, 2B). Net CO₂ assimilation remained below the compensation point under both 0.3 and 10 µmol m⁻² s⁻¹ green light (Fig. S1A). However, unlike for 10 µmol m⁻² s⁻¹, 0.3 µmol m⁻² s⁻¹ green light was not significantly different from dark-adapted conditions for stomatal conductance nor apparent electron transport rate (Fig. S1B-C). Together, these results indicate that dimG does not elicit detectable assimilation, electron transport or stomatal activity and is therefore physiologically equivalent to darkness.

**Figure 2.**
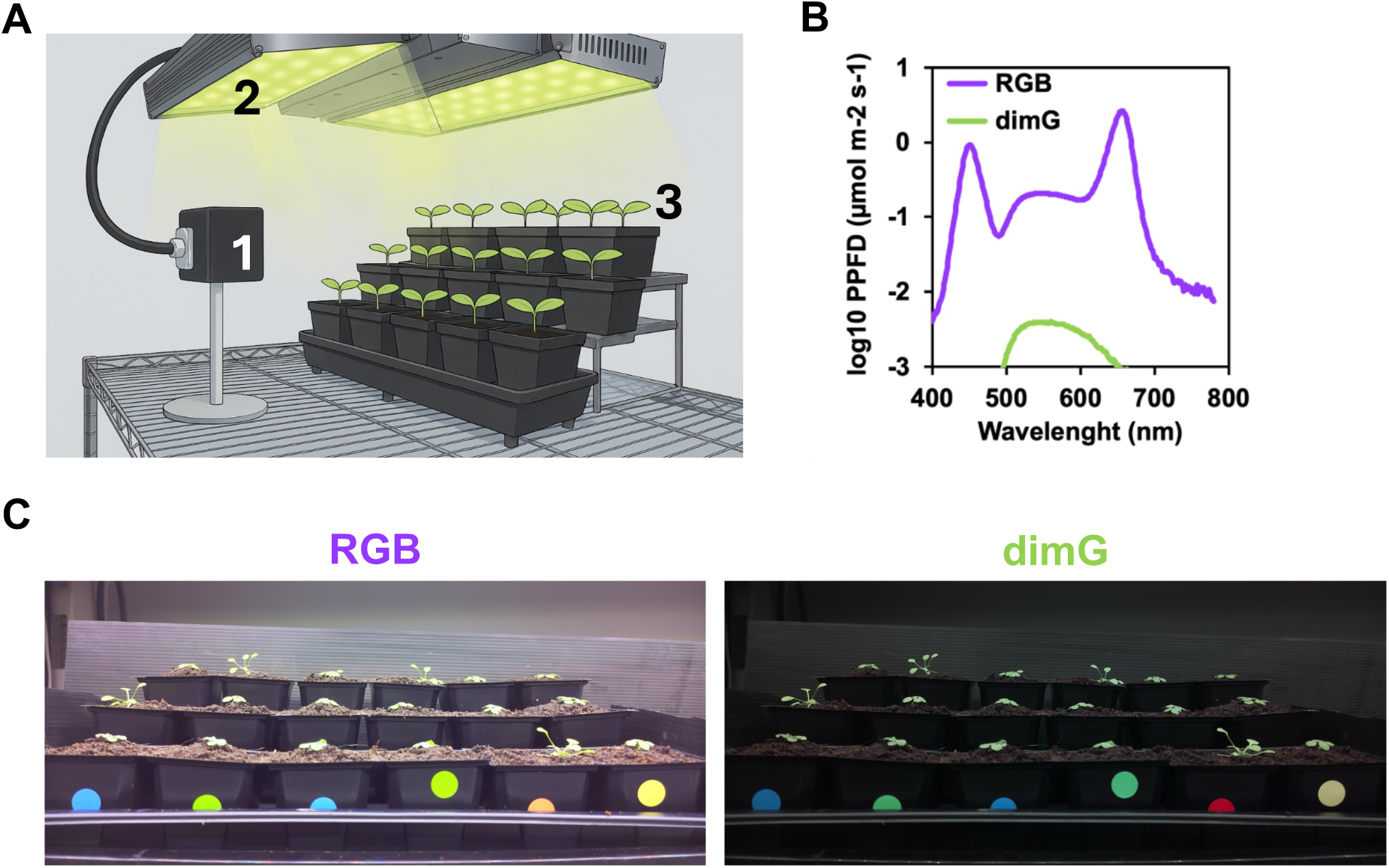
Imaging System for RGB and dimG acquisition. **(A)** Overview of imaging setup for *Arabidopsis* leaf movement analysis. (1) The Raspberry Pi Camera was mounted on a phone stand facing the plants; (2) Commercially developed horticultural LED fixtures (Vertical Future, UK) provided both RGB and dimG illumination; (3) Plants were arranged in three tiered rows to increase experimental capacity, and genotypes were distributed across rows to minimise potential effects of differences in light intensity. **(B)** Emission spectra used to support plant growth (RGB) and for imaging (RGB and dimG). **(C)** Representative images of *Arabidopsis* plants imaged under RGB or dimG illumination. A black panel was attached to the top row to provide a black background.

### Low-cost Raspberry Pi–based imaging system for RGB and dimG acquisition

We implemented a simple and cost-effective imaging setup using a Raspberry Pi Camera Module 3 NoIR (Fig. 2A, see Methods), enabling image acquisition over several days under full-spectrum RGB illumination during daytime and very low-fluence green light (0.3 µmol m⁻² s⁻¹, dimG) during nighttime (Fig. 2B-C). In addition, daytime dimG images were obtained by timing image acquisition to coincide with the dimG pulses from the illumination LEDs (Fig. 2B). To minimise noise arising from illumination fluctuations and leaf movements caused by airflow from the LED-panel fans, each final image corresponds to the average of 90 consecutive frames captured at 1 frame per second.

### Diel oscillations of leaf movement are altered in circadian oscillator mutants

Circadian rhythms are internal biological oscillations that allow plants to coordinate growth and physiology with the daily cycles in environmental conditions such as light and darkness (Harmer, 2025). In *Arabidopsis*, one of the most visible circadian outputs is the rhythmic upward and downward movement of leaves. The amplitude of this rhythm describes how pronounced the leaf movements are, whereas the phase indicates the timing of key events such as the peak or trough in leaf elevation. Hence, monitoring circadian rhythms of leaf movement under continuous light is a standard method in circadian biology for assessing circadian oscillator function (Edwards & Millar, 2007; Lou et al., 2022). In contrast, measuring leaf movements in light and dark cycles, which more closely resemble natural or controlled growth environments, is far less common (Bours et al., 2012; Dornbusch et al., 2014; Oskam et al., 2024). We tested whether dimG could support imaging suitable for quantifying diel leaf-movement rhythms. We examined how mutations that alter circadian period and rhythmicity influence diel leaf movement. Specifically, we compared wild type *Arabidopsis* (Col-0) with three well-characterized circadian oscillator mutants in a Col-0 background that have distinct circadian phenotypes under continuous free-running conditions: a short circadian period *light-regulated wd1* and *2* double mutant (*lwd1 lwd2*) (Airoldi et al., 2019), a long circadian period *zeitlupe (ztl-3*) mutant (Kevei et al., 2006), and a circadian arrhythmic *early flowering 3* (*elf3-1*) mutant (Hicks et al., 1996). These different plant genotypes were grown under full RGB during the day and under dimG at night.

We tested two imaging illumination regimes to capture the image sequence. In the RGB^Day^_dimG^Night^ regime, plants were imaged under RGB light during the day and dimG at night (Fig. 3A). In the dimG^Day_Night^ regime, daytime RGB illumination was briefly interrupted by pulses of dimG for image capture and nighttime images were also acquired under dimG, (Fig. 3A). We used the Tracking Rhythms in Plants (TRiP) algorithm to quantify leaf motion of whole-plants (Greenham et al., 2015). Under dimG^Day_Night^ imaging rhythms of plant leaf motion were detected in most plants, with *lwd1 lwd2* and *elf3-1* having higher amplitude oscillations than wild type and *ztl-3* (Fig. S2). Diel motion in wild type and *ztl-3* plants was broadly similar (Fig. S2A). In *lwd1 lwd2*, downward leaf motion extended during the day, while at night both *lwd1 lwd2* and *elf3-1* had prolonged upward movement (Fig. S2B-C). These data demonstrate that leaf movement is coregulated by light and the circadian oscillator, and disrupting circadian rhythms reveals the acute responses to light under diel cycles.

**Figure 3.**
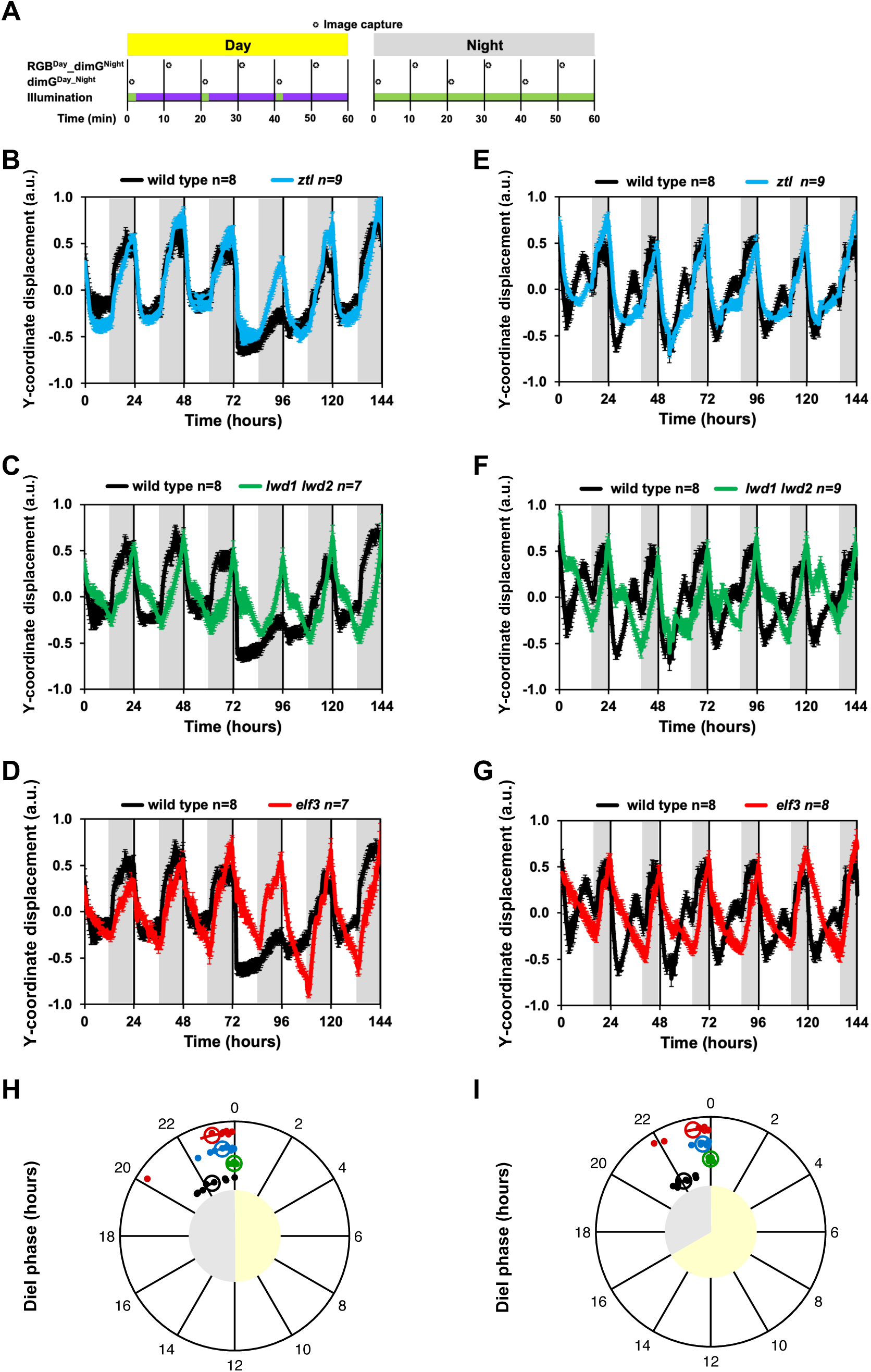
*Arabidopsis* circadian oscillator period and arrhythmic mutants have altered diel leaf movement rhythms. **(A)** Imaging schedule. Plants were imaged every 20 minutes. During the daytime, full-spectrum RGB illumination was used (indicated by a purple line) or dimG (indicated by a green line); during the night dimG illumination was used (indicated by a green line). Diel oscillations of plant vertical centroid position under neutral day (ND) (12h day, 12h night) and long day (LD) (16h day, 8h night) in *ztl-3* **(B, E),** *lwd1 lwd2* **(C, F),** and *elf3-1* **(D-G)** compared to wild type **(B-G).** The image sequence used for analysis was dimG^Day_Night^. Plots depict mean and SEM of the indicated number of plants (n). In B-F, day and night are indicated by white and grey shading, respectively **(H, I)** Diel phase of maximum vertical centroid position under ND (H) and LD (I). Circular plots show the average diel phase calculated over five consecutive 24-hour windows. Unfilled circles represent the genotype average diel phase, with lines indicating the SEM. Filled small dots represent the average diel phase of individual plants. Colours denote genotypes: wild type (black), *ztl-3* (blue), *lwd1 lwd2* (green), and *elf3-1* (red). Light yellow and light grey shading indicate day and night, respectively. Individual Y-coordinate displacement curves are shown in Fig. S4 (ND) and Fig. S5 (LD).

We tested whether the consistent illumination used in the dimG^Day_Night^ imaging regime was necessary for accurately quantifying leaf-movement dynamics. To do this, we compared the TRiP leaf-motion outputs from dimG^Day_Night^ with those from RGB^Day^_dimG^Night^, in which images were captured under RGB during the day and dimG at night. In plants with high-amplitude oscillations, leaf-motion outputs were highly similar between the two imaging regimes (Fig. S3A-D). In contrast, in plants with low amplitude oscillations, TRiP detected additional motion associated with the transitions between RGB and dimG illumination in the RGB^Day^_dimG^Night^ sequence (Fig. S3A-D). These results indicate that consistent illumination across the full-time image sequence, as provided by the dimG^Day_Night^ regime, yields more accurate leaf-motion outputs.

Because TRiP reports motion flux rather than absolute leaf position, it cannot directly identify the timing of maxima or minima in leaf elevation, which is necessary for determining phase. To examine how circadian and light cues establish the phase of leaf movement under diel conditions, we calculated the Y-coordinate displacement of the plant segmented silhouette (see Methods) from dimG^Day_Night^ image sequences to get an accurate measure of leaf positions over time under neutral (ND, 12 h light/12 h dark) and long-day (LD, 16 h light/8 h dark) photoperiods (Fig. 3). Under neutral days, wild type plants had sharp downward and upward movement at dawn and dusk, respectively, followed by stable positions (Fig. 3B-D). *ztl-3* had a similar profile to the wild type but with a delayed peak (Fig. 3B). In *lwd1 lwd2* and *elf3-1*, dawn and dusk movements were less sharp and continued until the next light transition (Fig. 3C-D). Under long days, wild type had an initial daytime peak followed by a higher nighttime peak (Fig. 3E-G). In *lwd1 lwd2*, the daytime peak was advanced (Fig. 3F), whereas in *ztl-3* and *elf3-1*, the daytime peak was absent (Fig. 3E and G). Nighttime peaks were delayed in all mutants compared to wild type and the phase of peak maxima was largely unchanged between photoperiods (Fig. 3G-H). Overall, photoperiod altered wild type diel profiles, producing a second daytime peak under long days. The effect of *ztl*-3 was more pronounced under long days, while *lwd1 lwd2* and *elf3-1* remained dominated by light–dark transitions. Notably, *lwd1 lwd2* was photoperiod-sensitive, having an early daytime peak under long days, whereas *elf3-1* profiles were similar across photoperiods. We noted that, unlike TRiP motion estimation, Y-coordinate displacement was highly similar between the dimG^Day_Night^ and the RGB^Day^_dimG^Night^ imaging regimes (Fig. S4, S5). These results confirm that dimG imaging supports accurate detection of genotype-specific differences in diel leaf movement, validating its use for circadian phenotyping under realistic light-dark cycles, a task that is difficult to achieve with most current imaging approaches.

### dimG enables nighttime imaging of leaf movement in lettuce

Applying dimG illumination offers a low-cost alternative for monitoring plant growth in controlled environments. As a proof of concept, we tested this approach using a commercial lettuce variety in a laboratory facsimile of a commercial vertical farm (Fig. 4A). Our first objective was to determine whether dimG illumination could support nighttime time-lapse imaging for quantifying leaf movement in lettuce. Because infrared (IR) illumination combined with IR-sensitive cameras is routinely used for nighttime imaging (Olas et al., 2020), we directly compared IR and dimG as alternative light sources.

**Figure 4.**
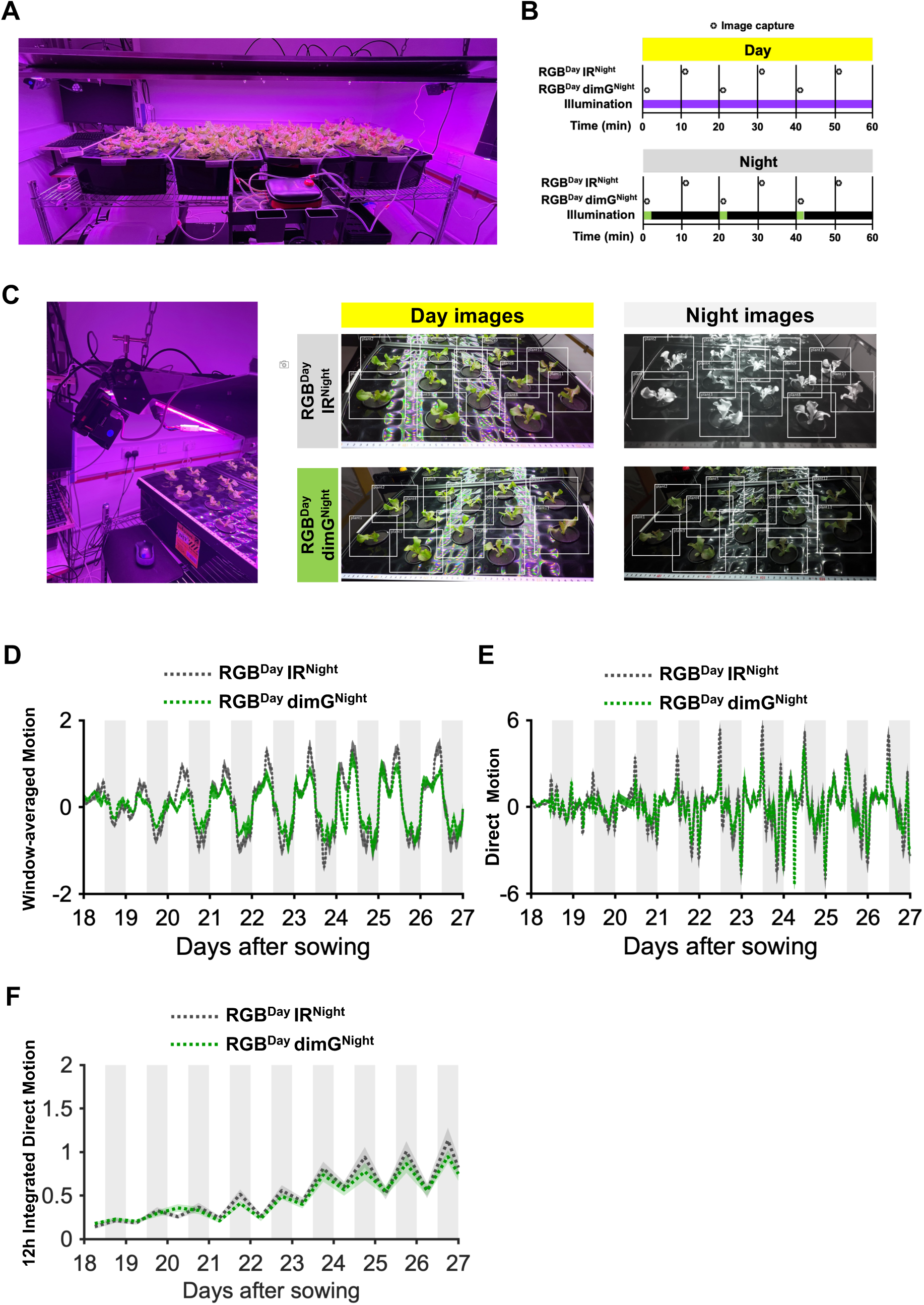
Plant leaf motion quantification is comparable when imaging under infrared (IR) or dim green (dimG) light during the night. **(A)** Overview of laboratory facsimile of a commercial vertical farm with lettuce grown hydroponically. **(B)** Imaging schedule. Plants were imaged every 20 minutes. During the daytime, full-spectrum RGB illumination was used (indicated by a purple line); during the night, imaging was performed using either IR (indicated by a black line) or dimG illumination (indicated by a green line). **(C)** Positioning of Raspberry Pi camera over hydroponic tank. Representative images from the RGB^Day^ IR^Night^ and RGB^Day^ dimG^Night^ image sequences. Boxes indicate the regions of interest (ROIs) used for TRiP-based leaf motion estimation. The same plants were imaged simultaneously under both nighttime conditions to enable direct comparison of leaf motion estimates (n=12). **(D)** Window-averaged Motion (dotted lines) ± SEM (shaded regions) for RGB^Day^ IR^Night^ (grey) and RGB^Day^ dimG^Night^ (green) conditions. **(E)** Direct motion (dotted lines) ± SEM (shaded regions) for RGB^Day^ IR^Night^ (grey) and RGB^Day^ dimG^Night^ (green) conditions. **(F)** 12h integrated motion (dotted lines) ± SEM (shaded regions) for RGB^Day^ IR^Night^ (grey) and RGB^Day^ dimG^Night^ (green) conditions. White and grey shaded in E-F panels indicate day and night, respectively. Data represent the means of 12 plants; error bars indicate SEM. A replicate dataset of this experiment can be found in Fig. S6. Individual plant plots for window-averaged motion and direct motion are shown in Fig. S7 and S8, respectively.

Lettuce plants were grown hydroponically under 12 h light/12 h dark cycles and imaged every 20 minutes (Fig. 4A). During the day, images were captured under RGB and during night, images were captured under IR (RGB^Day^ IR^Night^) or under dimG (RGB^Day^ dimG^Night^). For dimG imaging, a 2-minute dimG light pulse (0.3 µmol m⁻² s⁻¹) was synchronized with image acquisition (Fig. 4B). The camera was positioned above the hydroponic tray at a 45° angle to maximize capture of leaf movement (Fig. 4C). Leaf movement dynamics were analyzed using TRiP, which has been applied previously to estimate circadian oscillations in lettuce (Anton-Sales et al., 2025). We first used TRiP’s default setting, which applies a 13-frame moving average, and refer to this metric as *Window-averaged Motion*. Nighttime imaging under IR or dimG produced similar motion curves although RGB^Day^ IR^Night^ imaging produced increased diel maximum and minimum values (Fig. 4D). To capture all movements and detect potential artifacts, we removed the moving-average step and refer to this metric as *Direct Motion*. Direct Motion curves were comparable between RGB^Day^ IR^Night^ and RGB^Day^ dimG^Night^ image sequences although motion peaks at the day-night transitions were higher in the RGB^Day^ IR^Night^ (Fig. 4E). Motion output was quantified by integrating the area under the curve over 12-hour day or night intervals (*12h Integrated Direct Motion*), confirming similar integrated motion values for both imaging conditions. Notably, integrated motion was lower during the day than at night, with an increasing trend over the course of the experiment (Fig. 4F). These results demonstrate that dimG illumination provides a practical alternative to IR for nighttime imaging, enabling quantification of diel leaf movement dynamics in lettuce.

Continuous dim green (dimG) illumination during the night could provide a more practical solution for automated time-lapse imaging compared to programming intermittent pulses. However, before adopting this approach, it is essential to determine whether nighttime dimG affects diel physiological processes. Similarly to Arabidopsis, exposure to dimG did not elicit photosynthetic activity above the compensation point or stomatal opening in lettuce, indicating that dimG is also physiologically equivalent to darkness in this species (Fig. S1 D-F).

To further confirm the minimal effects of dimG on physiology, we performed two independent experiments comparing plants exposed to complete darkness or dimG during the night, and quantified leaf movement dynamics and overall growth as indicators of circadian and developmental integrity (Fig. 5). Direct motion was comparable between lettuce plants grown with darkness (RGB^Day^ IR^Night^) or dimG (RGB^Day^ dimG^Night^) during the night (Fig. 5A), with both conditions having increased motion amplitude over time (Fig. 5B). Differences in motion direction during early transitions likely reflect imaging artifacts caused by switching from RGB daytime images to nighttime images with lower pixel intensity (dimG) or monochrome (IR) (Fig. 5B). Importantly, 12h integrated direct motion was comparable between light regimes, particularly during the night, and the trend of higher nighttime motion was maintained, indicating that continuous dimG exposure does not significantly alter leaf movement (Fig. 5C). Across the experiment, the 12h integrated motion was consistently higher at night (Fig. S9A,

**Figure 5.**
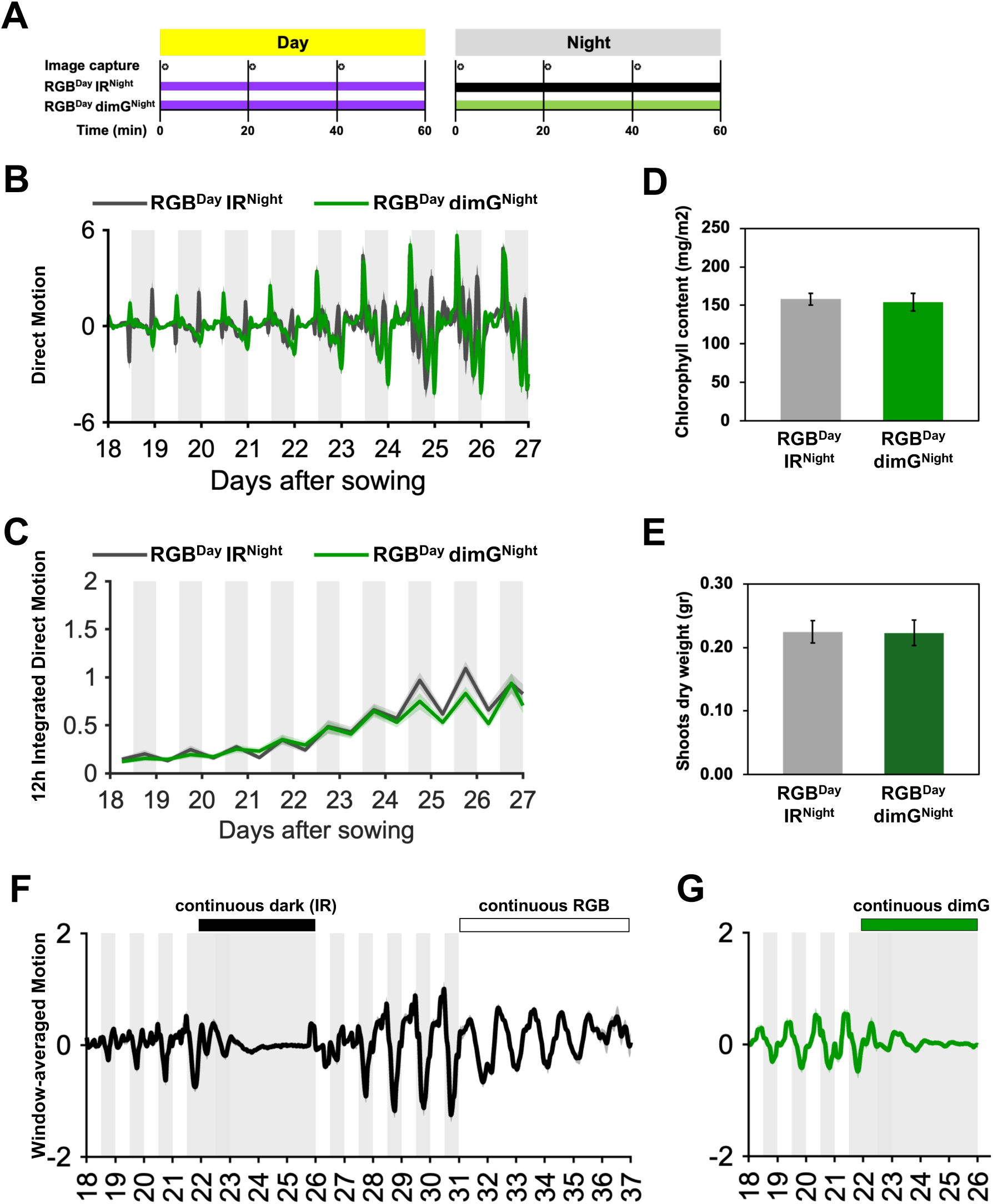
Nighttime exposure to continuous dimG does not influence leaf motion or overall plant growth in lettuce. **(A)** Imaging and lighting schedule. Plants were imaged every 20 minutes at the indicated times. During the day, full-spectrum RGB illumination was used (purple line); during the night, either infrared (IR; black line) or dim green (dimG; green line) illumination was applied. The photoperiod consisted of 12h day and 12h night. **(B)** Mean Plant Direct Motion + SEM for lettuce plants (n=12) grown under RGB^Day^ IR^Night^ (grey) and RGB^Day^ dimG^Night^ (green) conditions. White and grey shading indicate day and night periods, respectively. **(C)** Plant 12h Integrated Direct Motion(lines) ± SEM (shaded regions) for lettuce plants (n=12) grown under RGB^Day^ IR^Night^ (grey) and RGB^Day^ dimG^Night^ (green) conditions. White and grey shading indicate day and night, respectively. **(D-E)** Chlorophyll content (D) and shoot dry weight (E) of lettuce plants grown under RGB^Day^ IR^Night^ (grey) and RGB^Day^ dimG^Night^ (green) conditions, measured at day 28. Error bars indicate SEM **(F)** Continuous light, but not continuous darkness, sustains leaf movement oscillations in lettuce. Direct Motion grown under RGB^Day^ IR^Night^ (days 18-21 and 26-29), continuous darkness (IR; days 22-25) and continuous light (RGB; days 30-37). White and grey shading indicate light (RGB) and dark (IR) conditions, respectively. **(G)** Continuous dimG does not sustain leaf movement oscillations. Direct Motion for lettuce plants grown under RGB^Day^ dimG^Night^ (days 18-21) and continuous dimG (days 22-25). White and grey shading indicate light (RGB) and dark (dimG), respectively.

Wilcoxon signed-rank test, V=78 n=12, *p* < 0.001). To examine day-night differences in more detail, we calculated the average diel cycle of Direct Motion (Fig. S9B), which confirmed the similarity between the GB^Day^ IR^Night^ and dimG RGB^Day^ dimG^Night^ conditions. Daytime movement was largely confined to light-dark transitions, whereas multiple additional motion events occurred during the night (Fig. S9B). Total daily motion increased over the course of the experiment at a similar rate under both GB^Day^ IR^Night^ and dimG RGB^Day^ dimG^Night^ conditions (Fig. S9C). Consistently, plants exposed to dimG during the night had no differences in dry shoot weight or chlorophyll content at the end of the experiment (Fig. 5D-E). Overall, continuous nighttime dimG exposure does not alter plant leaf motion dynamics.

In *Arabidopsis*, prolonged darkness reduces leaf elongation and dampens leaf movement, whereas continuous light sustains circadian-driven oscillations (Dornbusch et al., 2014). We hypothesized that if dimG does not influence growth or circadian regulation, lettuce under continuous dimG would behave similarly to continuous darkness. Lettuce grown under diel cycles had robust rhythmic leaf movements (Fig. 5F, days 18–21) that were abolished when transitioned to continuous darkness; rhythms persisted for one day before damping (days 22–25) and were reinstated upon reintroduction of diel cycles (days 26–30). Under continuous light, oscillations persisted for at least five days (days 31–36). Similarly, under diel cycles of RGB and dimG, lettuce had rhythmic movements (Fig. 5G, days 18–21). However, under continuous dimG, rhythms persisted briefly before damping (days 22–25), mirroring the response to continuous darkness and confirming that dimG does not sustain circadian oscillations.

Taken together, these findings demonstrate that continuous dim green illumination enables high-frequency nighttime imaging without altering growth or circadian-regulated leaf movement, validating its use as a practical solution for automated phenotyping in laboratories, experimental growth facilities and controlled environment agriculture.

## Discussion

Our results demonstrate a practical and cost-effective strategy for continuous plant monitoring across diel cycles, with the potential to repurpose the green-light channel already present in many commercial LED growth panels for nighttime imaging. We show that extremely low-intensity green illumination (0.3 µmol m⁻² s⁻¹) is sufficient to capture high-quality images of both *Arabidopsis* and lettuce in a controlled environment representative of commercial vertical farming systems, without detectable disruption of circadian rhythms, or plant physiology. Low-cost image capture was possible because the Raspberry Pi Camera Module 3 used in this study contains an IMX708 sensor that is capable of low intensity imaging. At current prices, this camera alone is available for ∼US$25. This approach provides an accessible and easily deployable solution for continuous plant phenotyping.

We evaluated two methods for quantifying leaf-movement dynamics in our image sequences: TRiP (Greenham et al., 2015) and a colour-segmentation algorithm that estimates Y-coordinate displacement (see Methods). Both approaches require minimal manual segmentation, can be fully automated when plant positions remain constant across experiments, and run efficiently in Python. Importantly, they capture whole-plant movement rather than focusing on individual leaves (Bours et al., 2012; Dornbusch et al., 2014; Oskam et al., 2024). In Arabidopsis, we found that TRiP performs reliably under dimG illumination for both day and night imaging (dimG^Day_Night^) (Fig. S2, S3), whereas Y-coordinate displacement provides robust motion estimates when analysing both dimG^Day_Night^ and RGB-by-day and dimG-by-night image series (RGB^Day^ dimG^Night^) (Fig. 3, S4 and S5). Leaf movement has long been used to characterise circadian rhythm defects in mutants without the need for transgenic markers (Edwards & Millar, 2007). We demonstrate the applicability of our system under diel conditions by investigating how circadian oscillator mutations shape diel leaf-movement profiles. We found that *Arabidopsis* adjusts the diel movement pattern to the photoperiod, with wild type plants having a leaf-elevation peak at the end of the day followed by a higher night peak under long-day photoperiods (Fig. 3). A comparable double-peak profile under long-day photoperiods has been detected with near-infrared laser scanning (Dornbusch et al., 2014), showing that our approach can resolve leaf-movement dynamics previously accessible only through specialised techniques. Notably, although *elf3* mutants did not adjust their diel rhythms to photoperiod, mutants with altered circadian periods had photoperiod-specific responses (Fig. 3), highlighting extensive circadian control over diel leaf movement.

In agreement with a recent report (Anton-Sales et al., 2025), TRiP provides a reliable and accessible method for quantifying whole-plant leaf-movement dynamics over time in lettuce. We found that both diel and free-running leaf-movement dynamics can be effectively monitored using dimG illumination for nighttime imaging without affecting circadian rhythms (Fig. 4–5). By removing the window-averaging step from the TRiP workflow (see Methods), we obtained a direct motion estimate that preserved lower-frequency components of movement (Fig. 4E, 5B and S9B), revealing higher leaf motion during the night than during the day (Fig. 4F, 5C and S9A). A similar increase in nighttime motion was observed in *Arabidopsis* using the Phytotyping 4D platform which employs a light-field camera to generate three-dimensional reconstructions of plants (Apelt et al., 2015), consistent with our approach reliably capturing plant diel leaf-motion dynamics in lettuce. Furthermore, the progressive increase in directed motion as plants grew (Fig. 4D, 5C. S9C) suggests that direct motion captures growth-related components of leaf dynamics and may act as a proxy for plant growth, warranting further investigation in future studies. More broadly, our imaging approach using dimG could support monitoring tools for dynamic growth outputs, and plant development.

Taken together, this study demonstrates a low-cost solution for continuous quantification of diel plant leaf movements that mitigates technical and financial limitations of conventional high-resolution imaging platforms. By combining dimG nighttime illumination with accessible whole-plant motion-analysis methods, we provide a scalable approach for tracking diel and circadian dynamics in controlled environments.

## Methods

### Plant material and growth conditions

*Arabidopsis* genotypes used were *Col-0* (Airoldi et al., 2019), *ztl-3 (Haydon et al., 2017), lwd1 lwd2* (Airoldi et al., 2019) and *elf3-*1 (Herrero et al., 2025). *Arabidopsis* seeds were sown in individual 1 cm^3^ soil clusters (Levington M2 compost), stratified in the dark for 3 days at 4 ℃, and then placed under LED lights (conditions below). One week after germination, the soil clusters were transferred to 5 cm pot trays and grown for two additional weeks. At the start of the experiment, pots were saturated with water and arranged into tiered rows of six plants each (see Fig. 2), and plants were imaged for seven days under the conditions indicated in the figures.

Lettuce (*Lactuca sativa* var. *Outredgeous*) seeds (Sutton seeds, UK) were germinated in coco coir supplemented 5 mL/L of Ionic Hydro Grow (Growth Technology) dissolved in milliQ water and adjusted to pH 6 using pH UP (Growth Technology). Under LED lighting (conditions below, Fig. 4), seedlings were grown for 17 days before transfer to hydroponic culture. On day 17, seedlings were removed from the coco coir, washed with deionised water, placed into neoprene collars, and transferred to hydroponic baskets fitted into holes in the tank lid. Hydroponic tanks consisted of 30 L plastic tubs with lids containing twelve 5 cm-diameter holes, supporting twelve plants per tank. Each tank contained 24 L of hydroponic solution (10 mL/L of Ionic Grow pH 6 in milliQ water) with the following nutrient concentrations: nitrate 2.20% (w/v), phosphorus 0.35%, potassium 3.19%, calcium 1.16%, magnesium 0.44%, sulphur 0.12%, iron 0.04%, manganese 0.007%, boron 0.003%, zinc 0.003%, copper 0.002%, and molybdenum 0.001%. The solution was continuously aerated using an aquarium pump. The photoperiod was 12 h:12 h light:dark.

All leaf movement experiments were performed in a laboratory room with broad temperature control set to 20°C. Relative humidity was maintained at 50-60% using a digital hygrometer/humidifier (Govee H5075). Commercially developed horticultural LED fixtures (Vertical Future, UK), provided light conditions representative of those typically utilised in production settings. LED lights 30 cm above the plant canopy provided 160 µmol m^-2^ s^-1^ in the form of 105 µmol m^-2^ s^-1^ red (wavelength 600-700 nm), 20 µmol m^-2^ s^-1^ green (wavelength 500-600 nm), and 35 µmol m^-2^ s^-1^ blue (wavelength 400-500 nm) at plant height during the day (RGB). dimG, consisting of 0.3 µmol m^-2^ s^-1^ green (wavelength 500-600 nm), was provided during the night or in 2 min pulses coinciding with image capture as indicated in Fig. 3A, 4B and 5A.

### Leaf movement imaging system

The imaging system consisted of a Raspberry Pi Camera Module 3 NoIR (Raspberry Pi, SC0873), a Pi Compute Module 4 Rev5 (Raspberry Pi, CM4108016), and a Waveshare Mini Base Board Type B (Raspberry Pi, CM4-IO-BASE-B), all mounted in a 3D-printed enclosure. For *Arabidopsis* experiments, the imaging system was positioned to obtain a side view of the plants (Fig. 2). For the lettuce experiments, the cameras were positioned 20 cm above the plants at a 45° angle, centred along the longest side of each hydroponic tank (Fig. 4). Images were acquired every 20 minutes under either standard RGB illumination or dimG, as indicated in the figures. For imaging with dimG during the day or at night when indicated, a 2-minute dimG pulse was applied coinciding with image acquisition. Image acquisition was controlled with the Picamera2 library. Each image represented the average of 90 consecutive frames to minimise high-frequency leaf movements caused by airflow from the LED-panel fans and to reduce illumination variability during dimG pulses. Average images were saved as JPEG files at a resolution of 3840 × 2160 pixels in RGB format. Autofocus was enabled with a 2 sec settling period prior to acquisition for the *Arabidopsis* side imaging. Exposure and gain were controlled automatically unless otherwise specified.

### Quantification of leaf movement dynamics with TRiP

Using custom MATLAB scripts (The MathWorks Inc., version R2024b), missing frames were replaced with black images to prevent artificial gaps. For consistency in the analysis, the first image of the sequence corresponded to Time 0 (lights on) of the day after plants were transferred to the leaf movement stage for *Arabidopsis* or to the hydroponic tanks for lettuce. The MATLAB Image Segmenter app was used to pick a Region of Interest (ROI) wide enough to account for plant growth over the time of the experiment (see example in Fig. 4C). ROIs with no plants (and therefore no movement) were also selected to control for artificial background motion. ROI coordinates were extracted with a custom MATLAB script, and cropped image sequences were generated using the TRiP ‘CropAll’ function (Greenham et al., 2015). The motion within each crop image sequence was estimated with the TRiP ‘estimateAll’ function (Greenham et al., 2015). For *Arabidopsis*, the ‘estimateMotion’ script was used as originally intended with a motion output consisting of a moving average over 13-frames and we called it *TRiP Window-averaged Motion* (Greenham et al., 2015). In the lettuce experiments, we refer to the moving average over 13-frames as *Window-averaged Motion*. We also modified the ‘estimateMotion’ script to reduce the averaging window (line 83: taps = 13 replaced by taps = 1) and we called this output *Direct Motion*. We then estimated *the 12h-integrated direct motion* by computing the absolute trapezoidal (trapz) area under the curve for day or night interval with a custom script. For each tank, average values and standard error of the mean (SEM) were calculated.

*Diel direct motion* curves were calculated by converting experimental time to time-of-day and averaging replicate measurements at matching times across all days. Mean growth and SEM were then calculated for each time point.

### Quantification of leaf movement dynamics using Y-coordinate displacement

For Y-coordinate displacement analysis, we used a custom image-segmentation pipeline implemented in Python with OpenCV (Bradski, 2000). Seedling positions within each frame were defined once per experiment. Rectangular ROIs were drawn manually around each seedling of a representative frame. ROI coordinates and seedling IDs were then saved for cropping and subsequent analyses. Individual seedlings were cropped out of each frame within the time-series using a cropping script that applied every ROI to each frame. For each time point, the script extracted one crop per ROI and saved them into seedling-specific folders while preserving temporal order via filename. Overlay preview images showing ROIs on selected raw frames (i.e., first and last frame) were generated for visual verification.

Seedlings within each cropped image were segmented using a Python script implementing a multi-channel ensemble segmentation algorithm. Three complementary binary masks were generated: (1) an HSV-based “green” mask using adjustable hue (default 40–80°), saturation (≥50), and value (≥40) thresholds, cleaned with morphological opening and closing; (2) a CIELAB a-channel mask using inverted Otsu thresholding to detect green tissue, followed by morphological closing; and (3) a grayscale mask created by applying contrast-limited adaptive histogram equalization (CLAHE), Gaussian blurring, and Otsu thresholding. These three binary masks were then combined by equal-weight pixelwise voting (weight 1/3 per mask): for each pixel, we computed the mean of the three masks and classified it as seedling tissue if at least two of the three methods agreed (mean ≥ 2/3). The resulting consensus mask was morphologically cleaned (opening and closing) to remove small artefacts and fill gaps, yielding the final binary segmentation.

In the case where segmentation resulted in disconnected seedling fragments (such as when petioles were very thin), they were merged into a single segmented object representing the entire seedling. Segmented components (i.e. flecks of soil) below a minimum, pre-defined area were discarded as noise. For each time point, morphological parameters were calculated including area, perimeter, centroid, convex hull area, solidity, aspect ratio, circularity and number of merged components. Results were aggregated into a single CSV file. Optional outputs for visual verification included binary segmentation masks, coloured overlays on original images, side-by-side montages, and time-series grids showing seedling segmentations at regular intervals.

Y-coordinate displacement curves were linearly detrended and normalized to [-1,1] with BioDare2 (Zieliński et al., 2022). BioDare2 was also used to estimate free-running period and to identify non-rhythmic plants under diel cycles (real amplitude error < 0.5). Detrended and normalized curves were used to estimate phase of maximum Y-coordinate position for each 24 h window with a custom MATLAB script.

### Gas exchange measurements

Instantaneous gas exchange was measured in the dark and under 0.3 µmol m⁻² s⁻¹ green light, 10 µmol m⁻² s⁻¹ green light and full light conditions in both Arabidopsis and lettuce plants, after at least one hour under the light treatments. Both a LI-6800 infra-red gas analyzer and LI-600 porometer/fluorometer (LI-COR, USA) were used to measure assimilation, stomatal conductance, and apparent electron transport rate (aETR). For the LI-6800, the small light source with a clear top was used. Environmental conditions were as follows: CO2 concentration 400 ppm, 25°C, RH 60%, flow rate 500 µmol s⁻¹, fan speed 10,000 rpm, with no light provided by the instrument. Stability criteria were set at the default, with a wait time of approximately 4 minutes. Three (Arabidopsis) or five (lettuce) leaves on different plants were measured and assimilation and stomatal conductance determined. For the LI-600, stomatal conductance and aETR were measured, using the auto porometer and fluorometer setting, on one leaf of five different plants of each species. Two-way ANOVAs with Tukey’s multiple comparisons tests (alpha 0.05) were conducted, with species and light as the factors of interest.

### Chlorophyll content and weight measurements

Where indicated, lettuce plants were harvested at day 28 post sowing. Chlorophyll content was first measured using a chlorophyll meter (CCM-300, Opti-Sciences, US) at four positions per leaf on the third youngest leaf. Plants were then cut at the intersection between the shoot and root, with both parts patted dry and weighed for fresh weight before drying at 60 °C for at least 48 h. Shoots and roots were then weighed for dry weight.

### Bioluminescence quantification experiments

*pCCA1::LUC* luminescence assays were performed as previously described (Airoldi et al., 2019) with a few modifications. Plants were sown in clusters of 5-10 seeds and grown for 10 days under entraining conditions (12h-day, 12h-night, constant 20°C) in a growth cabinet under white light 100 µmol m^-2^ s^-1^. On day 10, plants were dosed with 4 mM D-Luciferin and transferred to the imaging system. Every hour, the bioluminescence signal was integrated for 800 seconds. Entrainment conditions were maintained for 2-3 days followed by 5 days of free-running conditions as indicated in Fig. 1. Luminescence intensity was extracted from each plant cluster using ImageJ. BioDare2 amplitude & baseline detrending (https://biodare2.ed.ac.uk/) was applied to the bioluminescence intensity curves and FFT-NLSS analysis from 24 - 120 h after continuous darkness or green light was performed to estimate the number of rhythmic plants (RAE <0.5) and the corresponding circadian periodicity (Zielinski et al., 2014; Zieliński et al., 2022).

## Data Availability Statement

The data supporting the findings of this study are available within the Supporting Information files accompanying this article. Image data generated during the study are available via the Adelaide University repository (https://doi.org/10.25909/32070474). Supporting datasets and scripts used for data processing and analysis are deposited in the University of Cambridge Apollo Repository (DOI: 10.17863/CAM.130057 and DOI: 10.17863/CAM.130058, respectively). Scripts for quantifying Y-coordinate displacement (ROI selection, image cropping, and ensemble segmentation) are deposited in the GitHub repository (https://github.com/DannyGinz/seedling-segmentation) and will be made publicly available upon publication.

## Acknowledgements

The authors acknowledge that this work was conceived during a collaboration funded by the United Kingdom Space Agency International Bilateral Fund and the Australian Space Agency Phase 2 grant 10089325. All authors acknowledge support from this grant, which included additional partners from Axiom Space, Saber Astronautics, and the University of Southern Queensland. In particular, we thank Cheryl McCarthy (University of South Queensland, Australia) for designing the code for image averaging. This work was also funded by the Australian Research Council Centre of Excellence in Plants for Space (CE230100015). We thank Wendy Sullivan and Charlotte Bampton (Adelaide University, Australia) for their expertise and plant growth support.

## Author contributions

Conceptualization: E.H, A.A., J.R.B., J.C.M., M.G., A.H.M, A.A.R.W. Methodology: E.H., D.N.G., A.R.G., A.A. Investigation: E.H., A.R.G., S.W., J.D.S. Formal analysis: E.H, A.R.G. Visualization: E.H. Writing - original draft: E.H. and A.A.R.W. Writing- review & editing: all authors. Supervision: J.R.B., J.C.M., M.G., A.H.M, A.A.R.W.. Funding acquisition: A.A., J.R.B., J.C.M., M.G., A.H.M, A.A.R.W.

## Supplementary Figures

**Figure S1.**
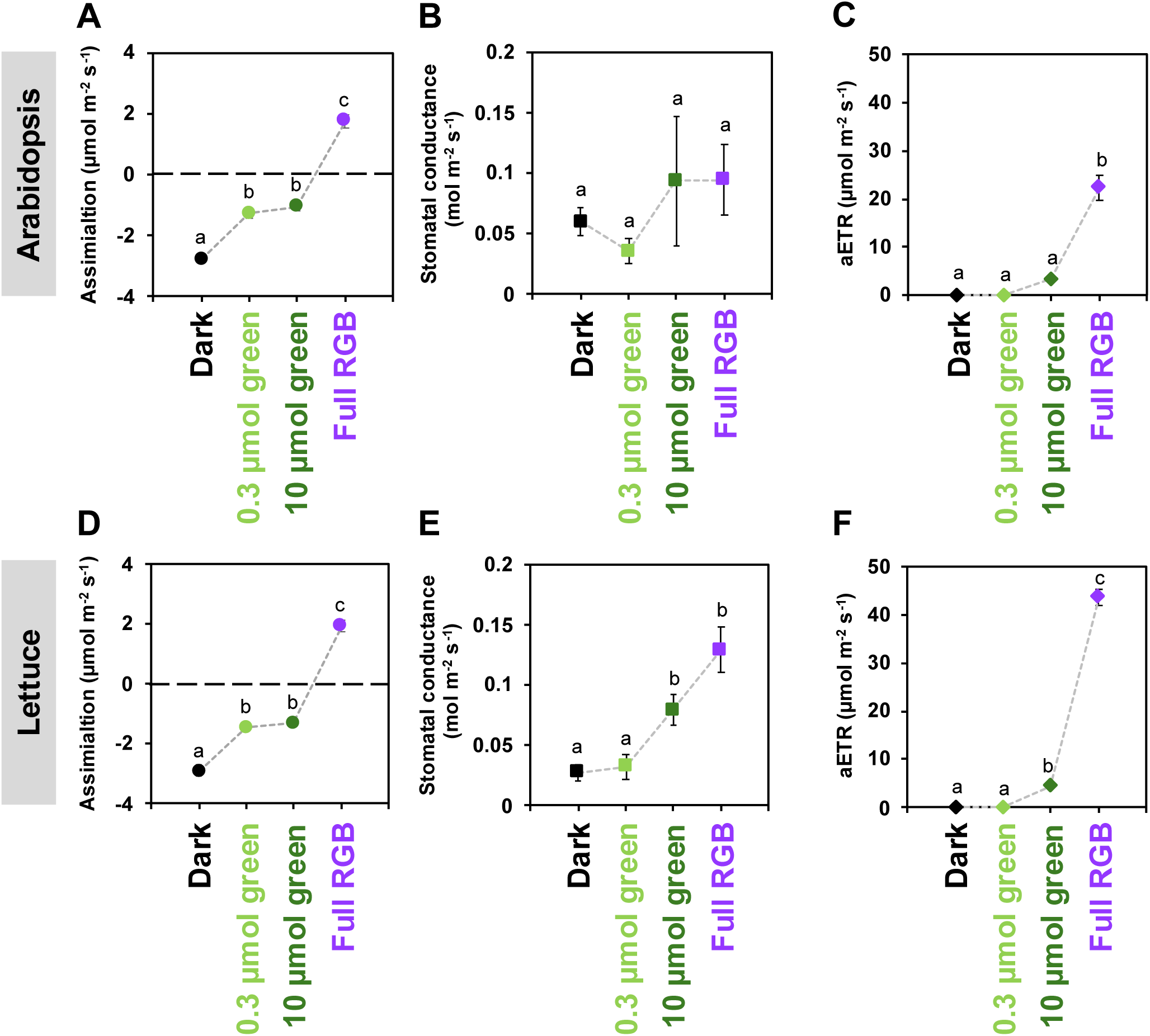
Dim green (dimG) light does not induce stomatal opening in Arabidopsis or lettuce, with minimal changes in assimilation and no electron transport. Assimilation in Arabidopsis **(A)** and lettuce **(D)**. Stomatal conductance in Arabidopsis **(B)** and lettuce **(E).** Apparent electron transport rate in Arabidopsis **(C)** and lettuce **(F)**. All measurements were taken after exposure to at least one hour of darkness, 0.3 µmol m⁻² s⁻¹ green (dimG), 10 µmol m⁻² s⁻¹ green, and full RGB light. Mean ± SEM (n = 3-5).

**Figure S2.**
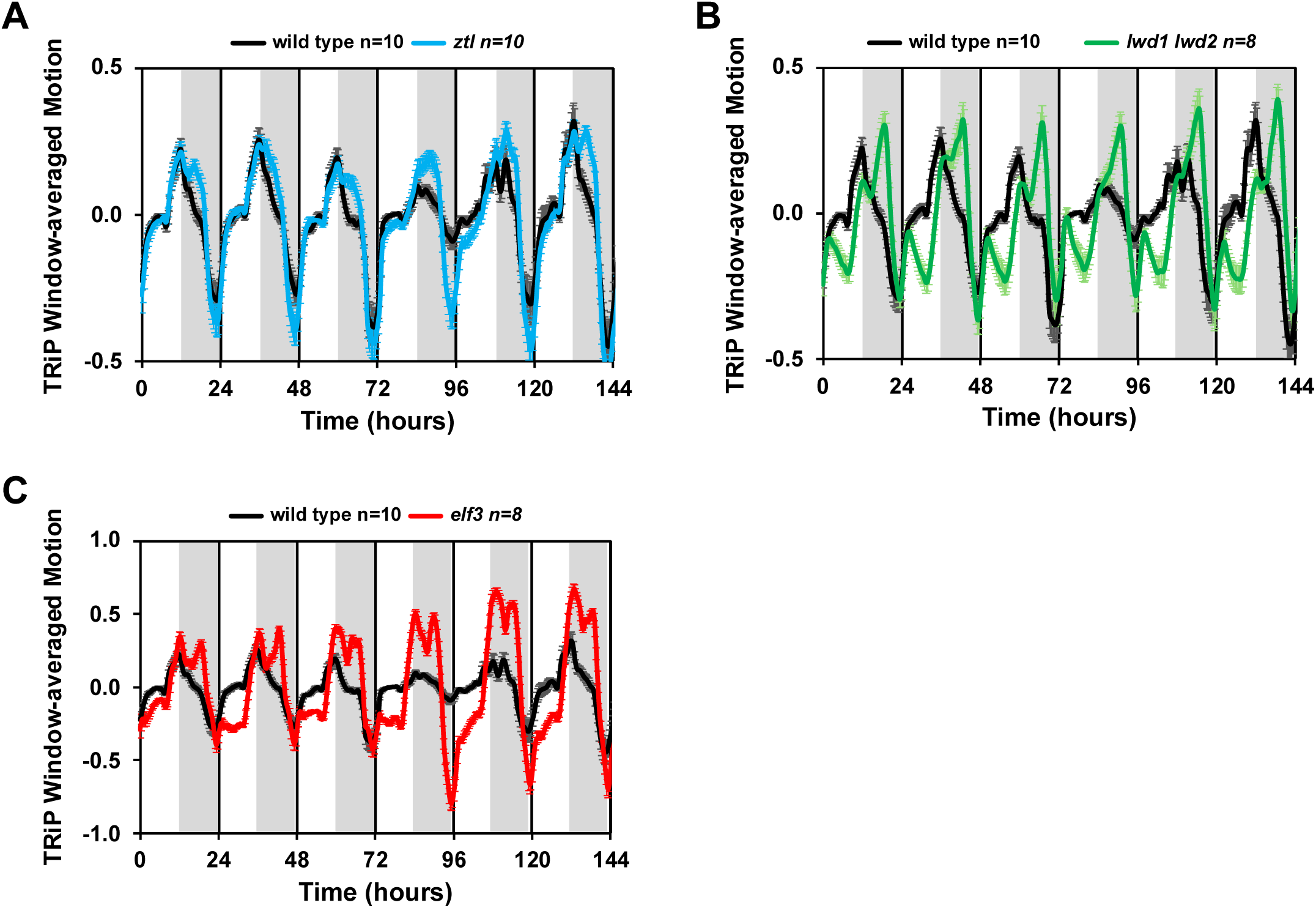
Diel leaf movement in *Arabidopsis* using dimG^Day_Night^ image sequence and TRiP. Genotype TRiP Window-averaged motion under ND (12h Day, 12h Night) **(A)** *ztl*, **(B)** *lwd1 lwd2*, and **(C)** *elf3* compared to wild type **(A-C)**. The image sequence used for the analysis was dimG^Day_Night^. Plots depict mean and SEM of the indicated number of plants (n). Day and night are indicated by white and grey shading, respectively

**Figure S3.**
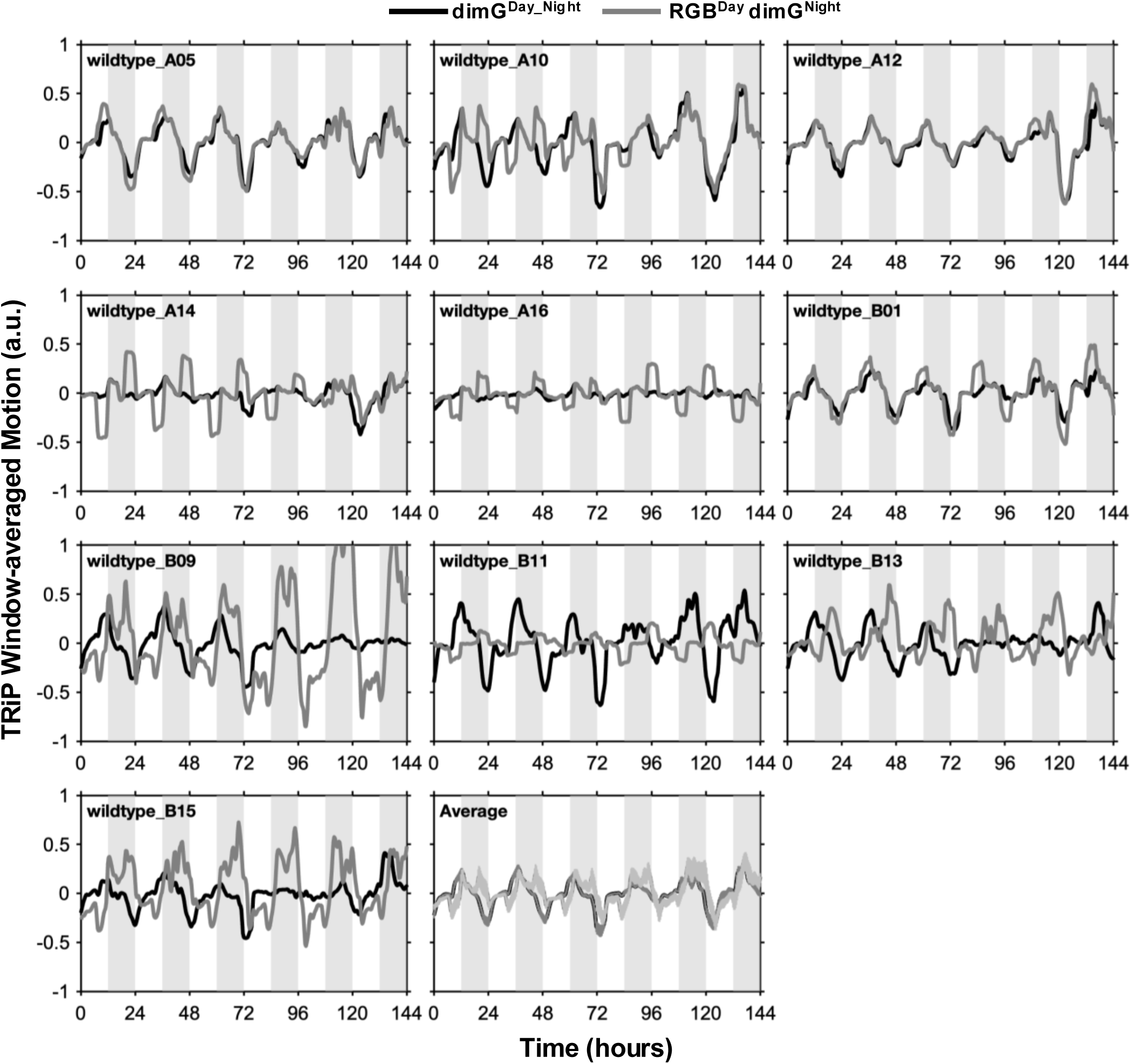
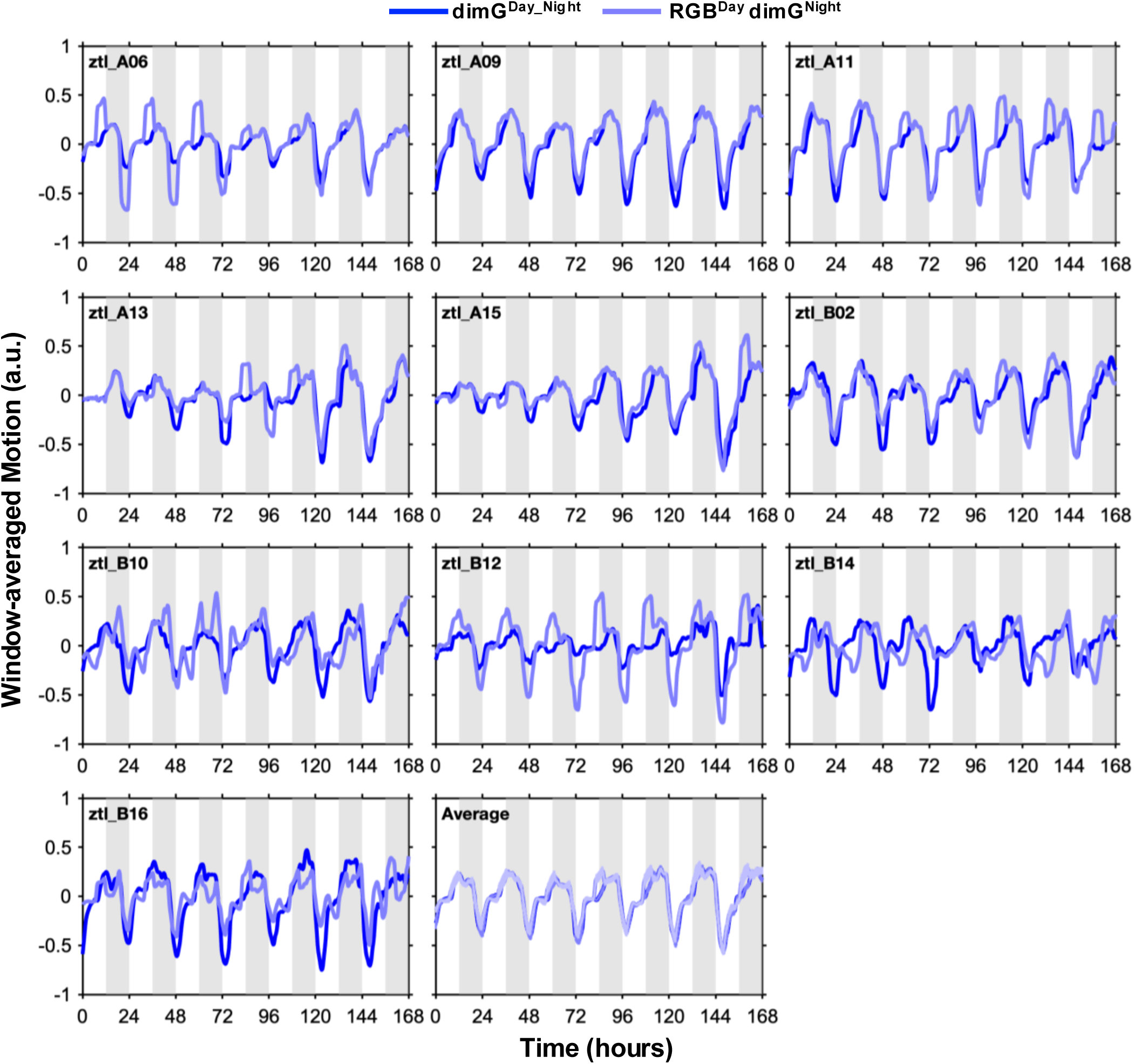
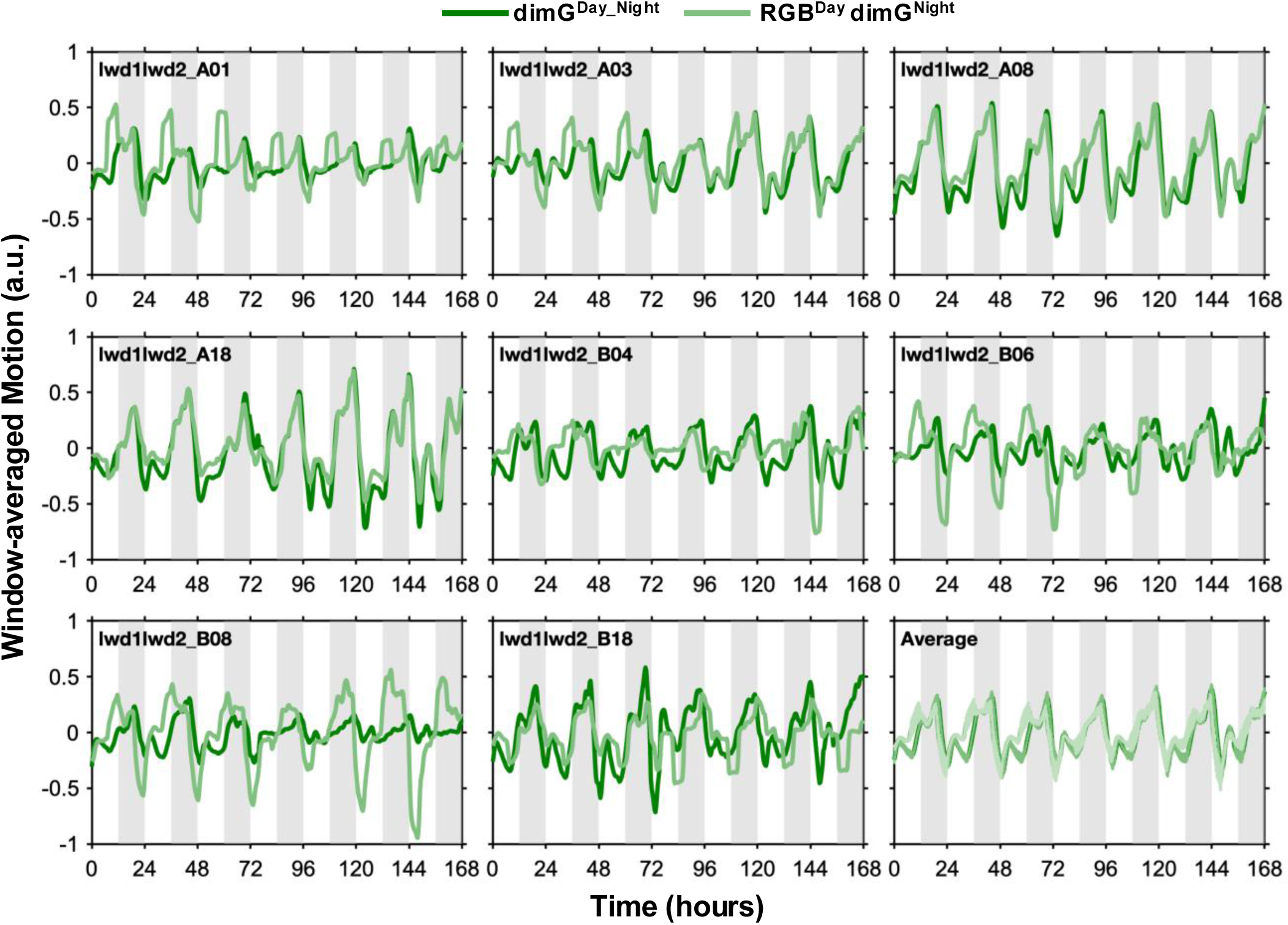
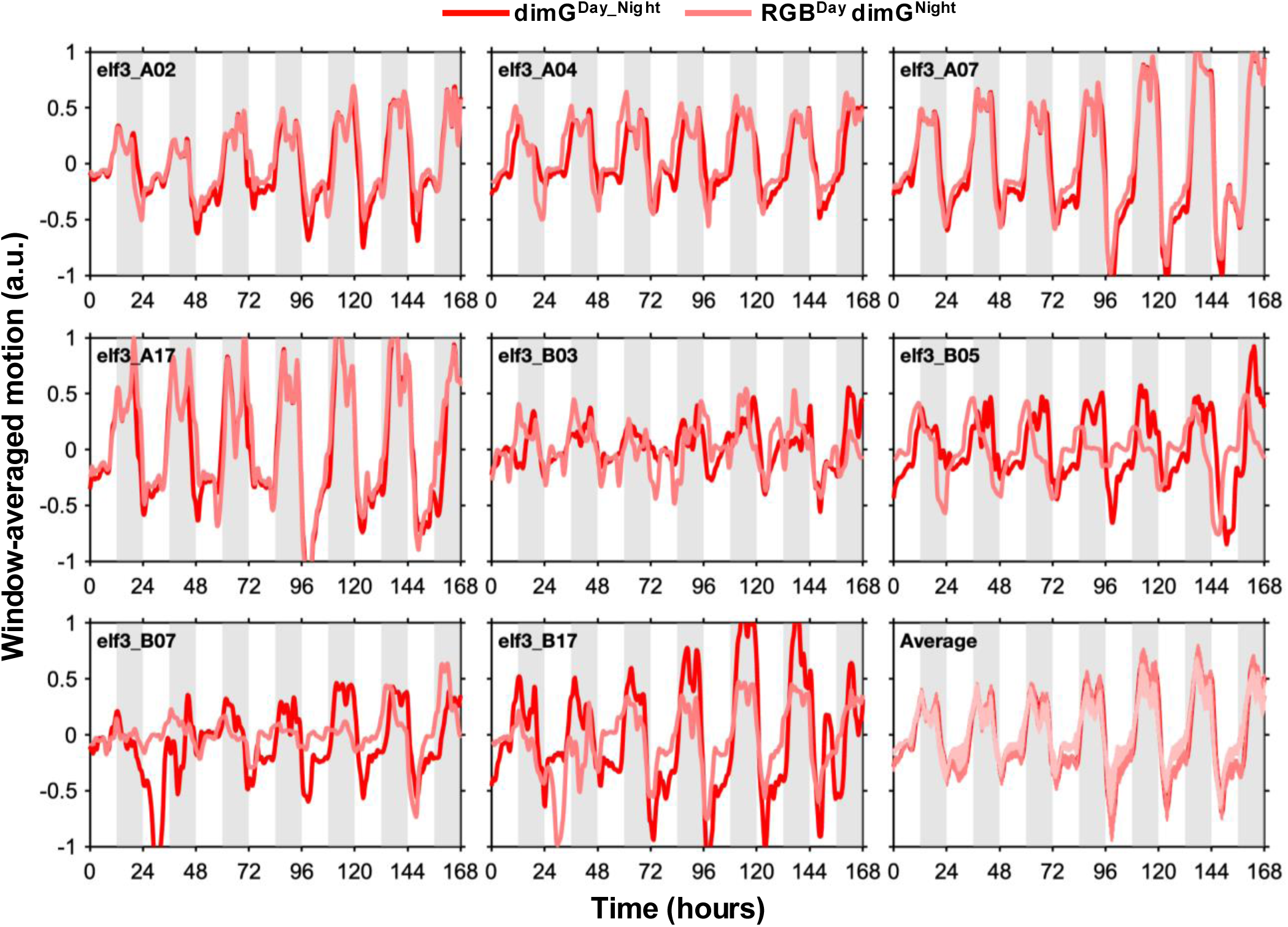
Window-averaged motion for individual plants using dimG^Day_Night^ and RGB^Day^_dimG^Night^ image sequences and TRiP. **(A)** Wild type, **(B)** *ztl*, **(C)** *lwd1 lwd2 and* **(D)** *elf3*.

**Figure S4.**
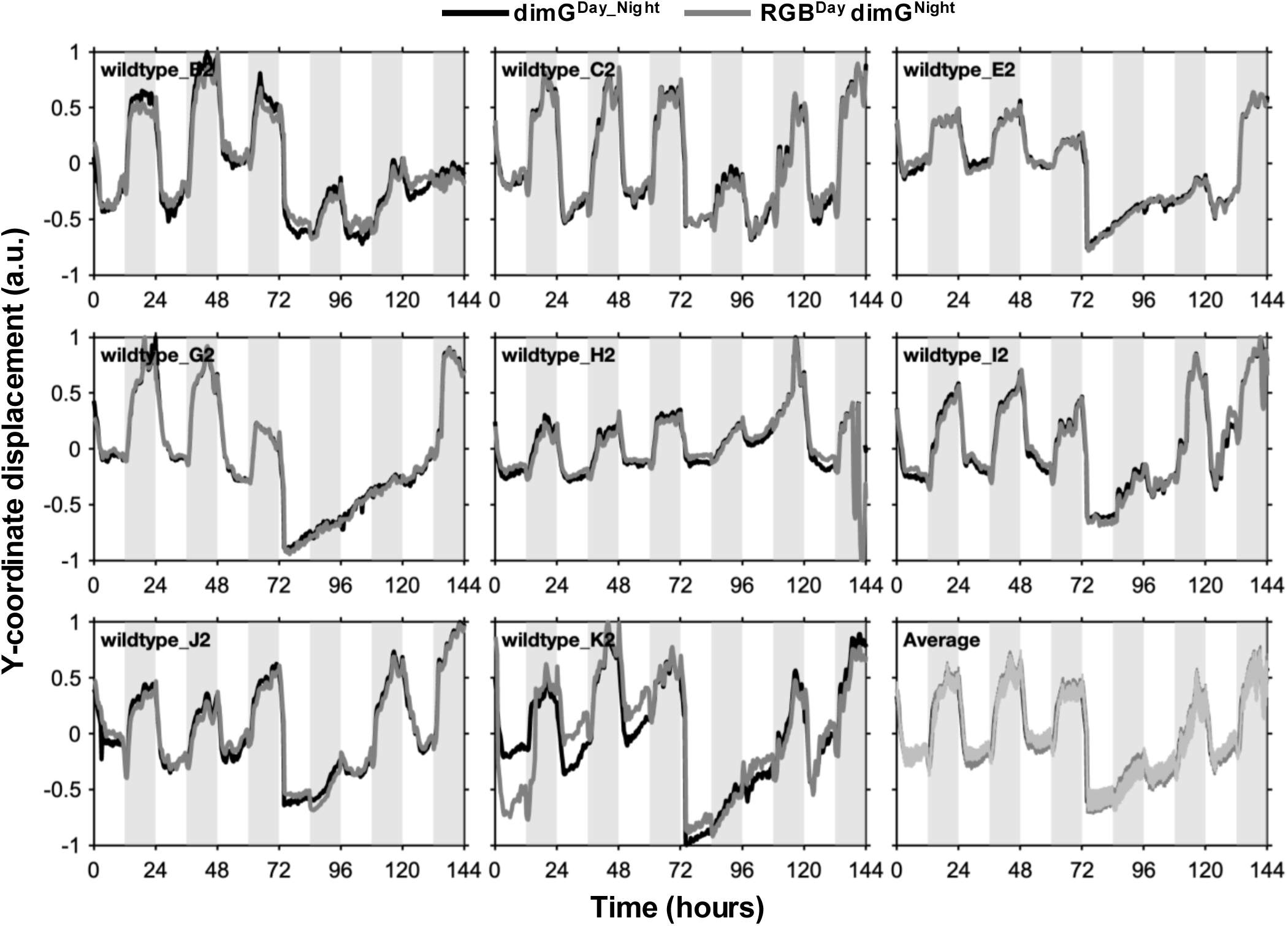
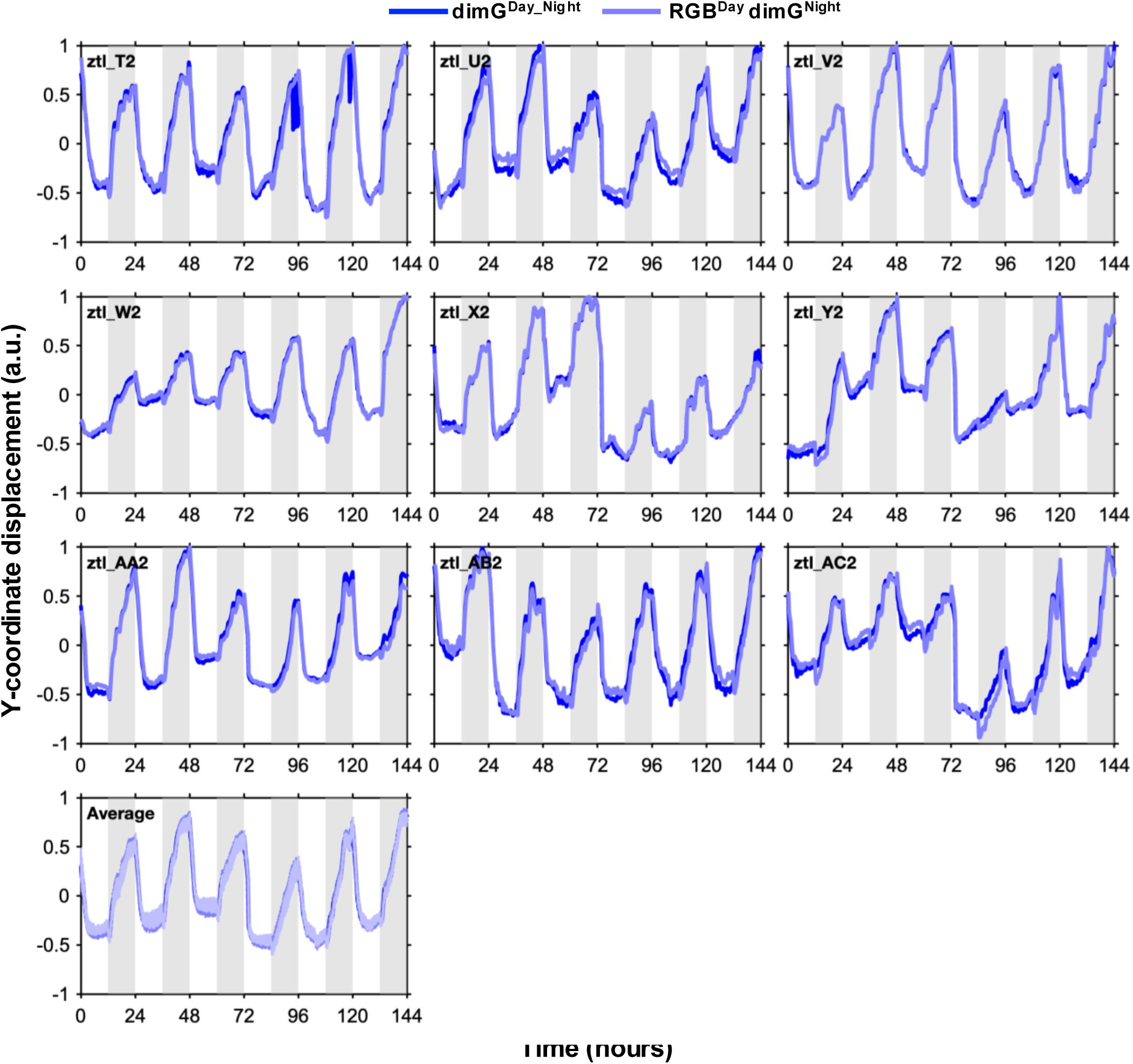
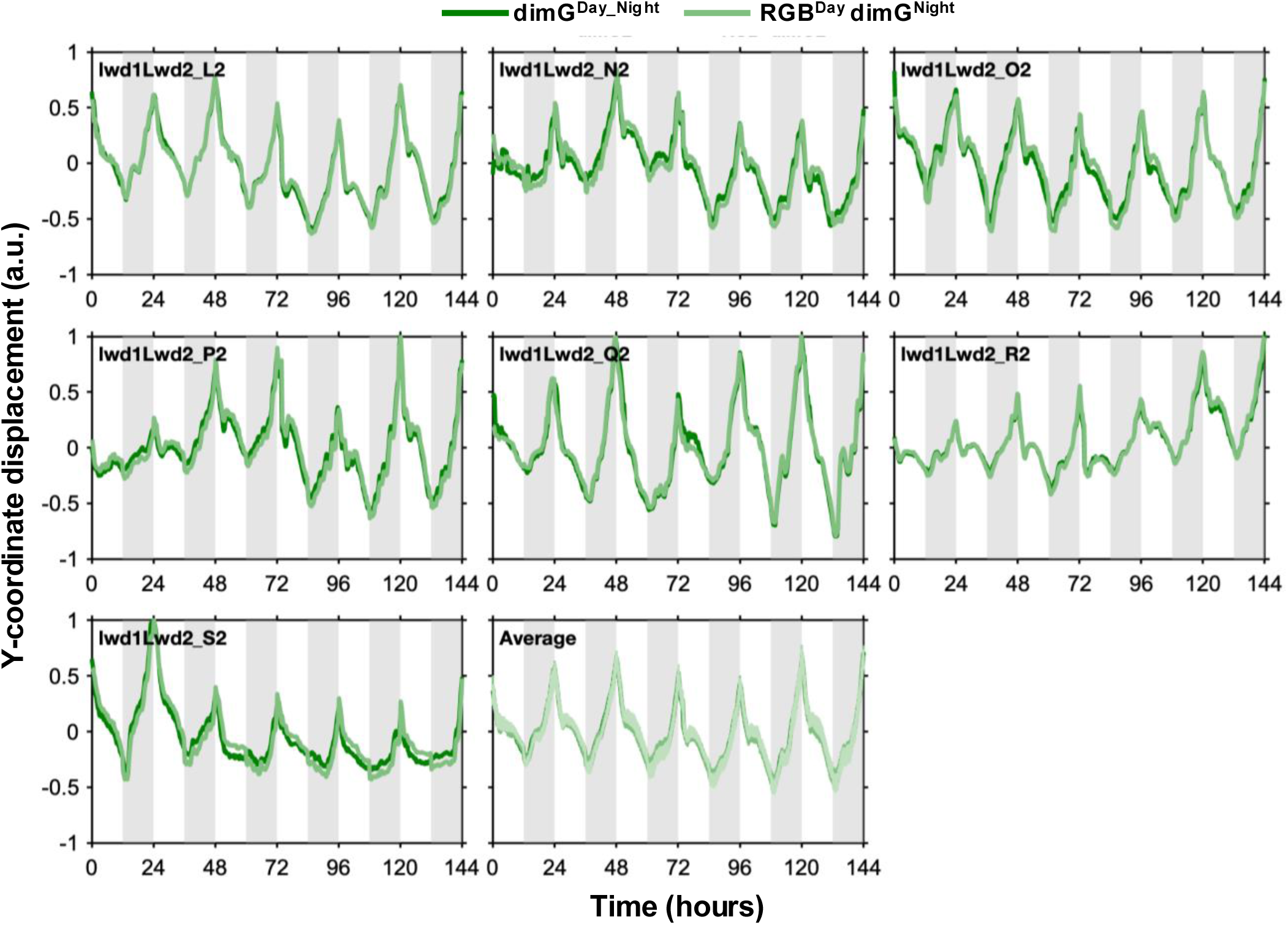
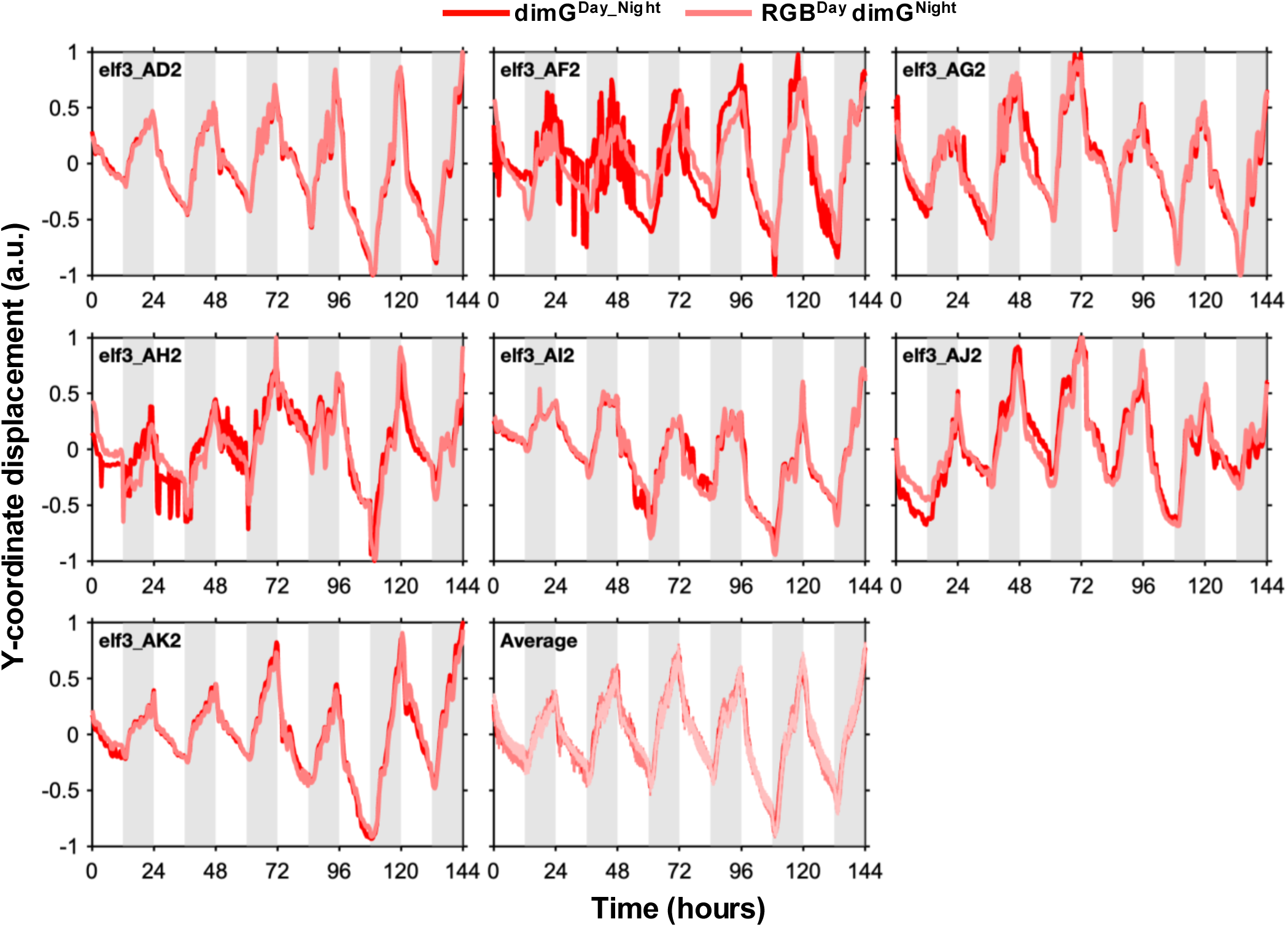
Diel oscillations of Y-coordinate displacement for individual plants using dimG^Day_Night^ and RGB^Day^_dimG^Night^ image sequences under ND. **(A)** Wild type, **(B)** *ztl*, **(C)** *lwd1 lwd2 and* **(D)** *elf3*.

**Figure S5.**
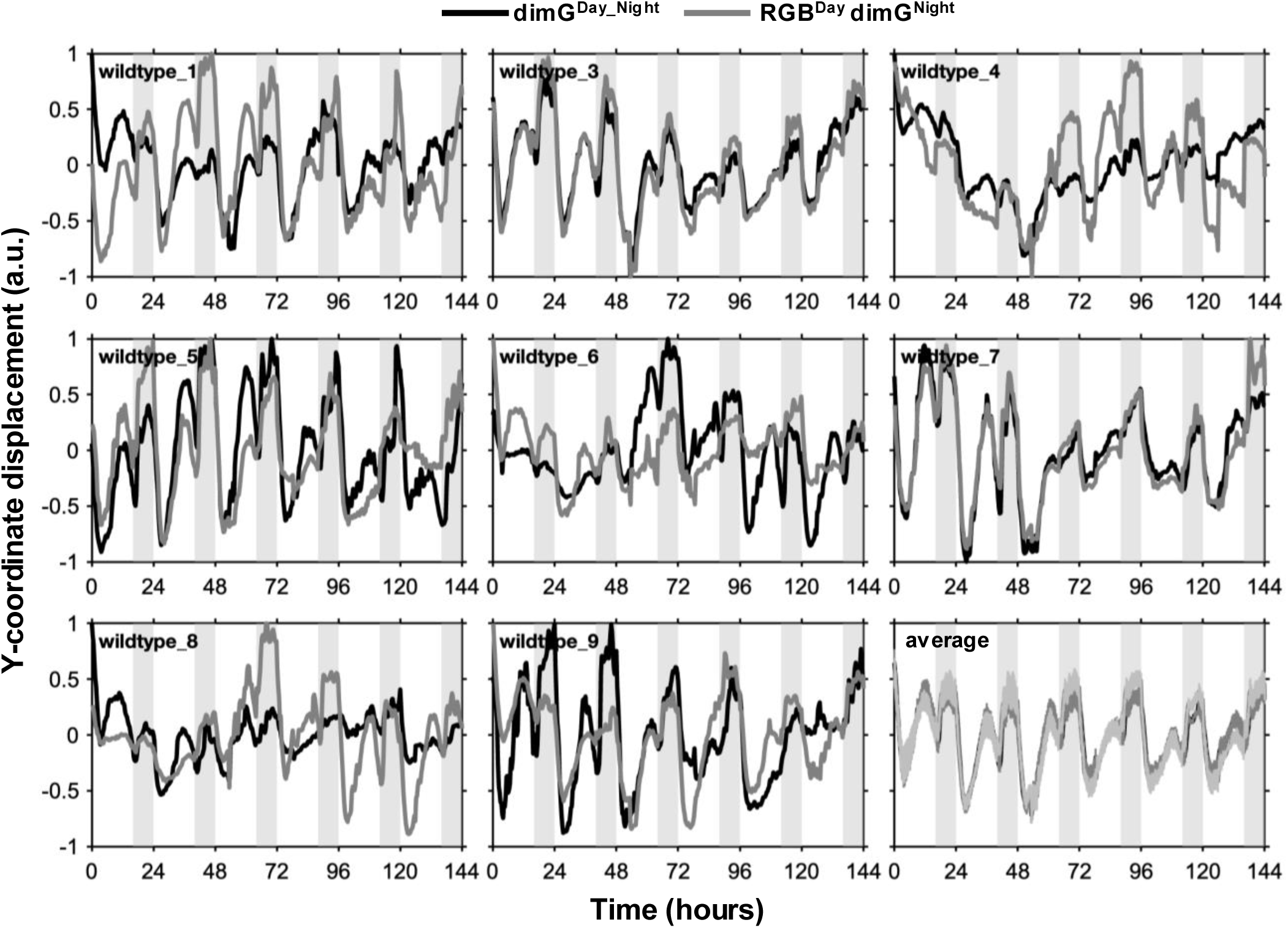
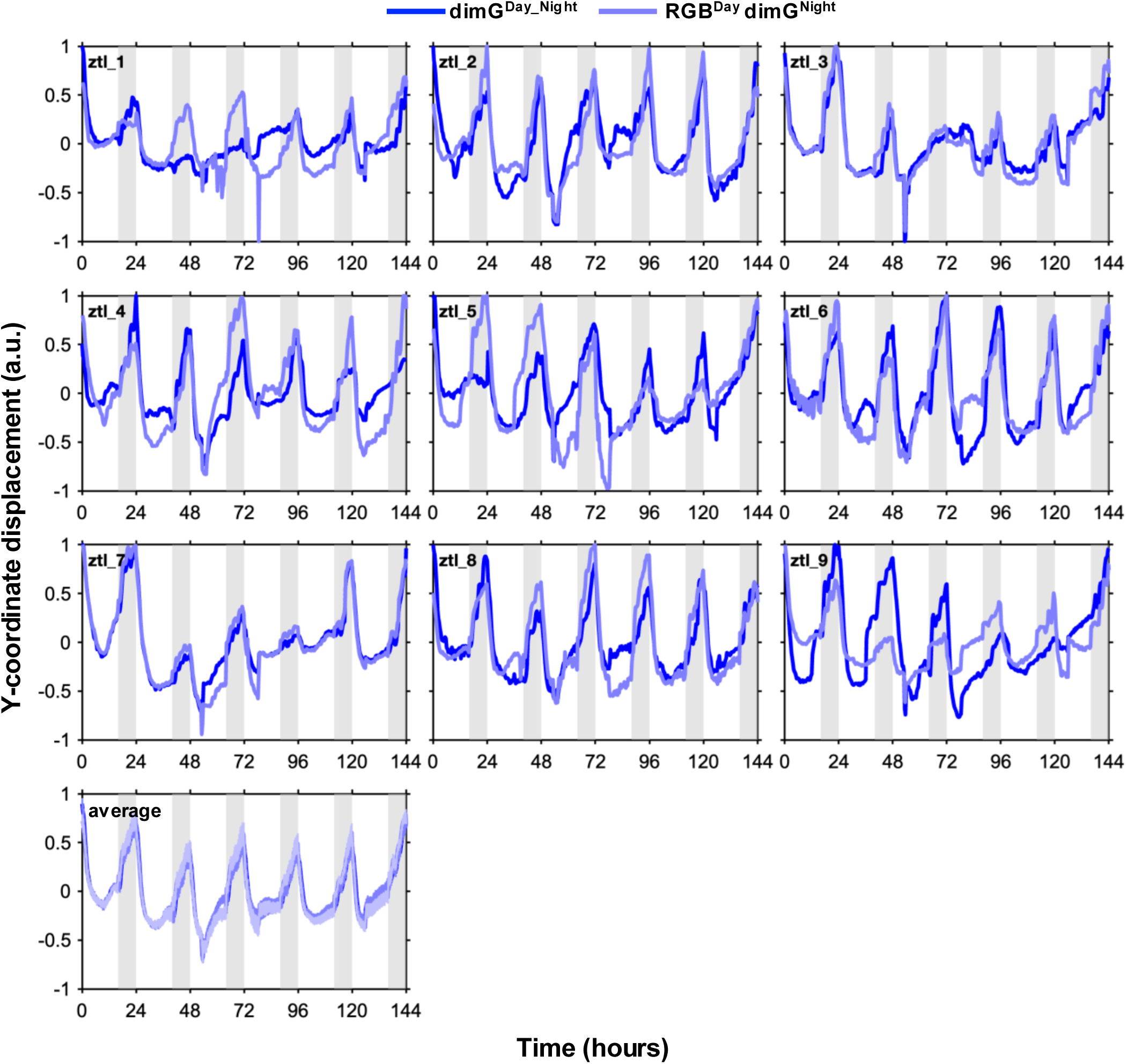
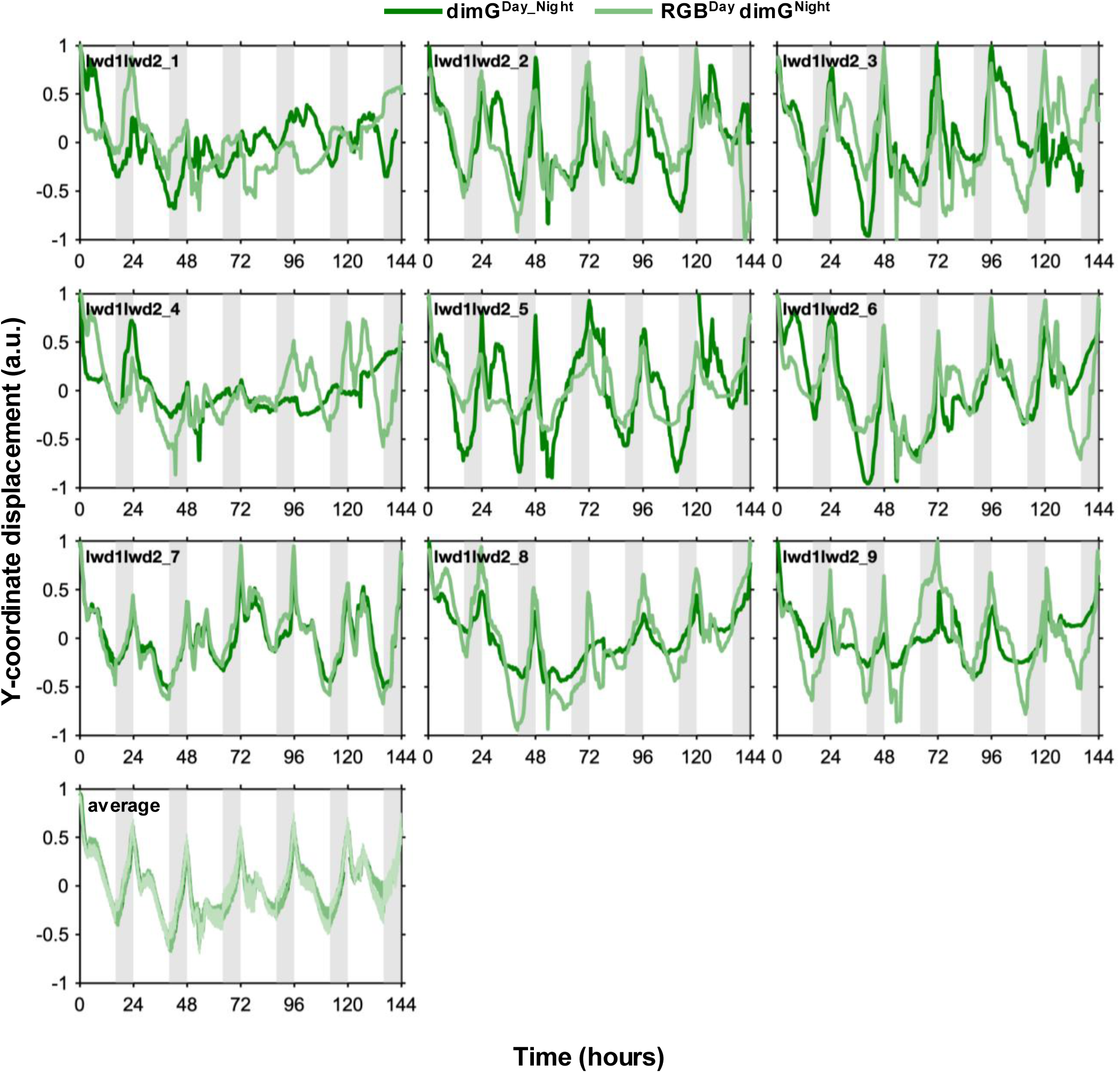
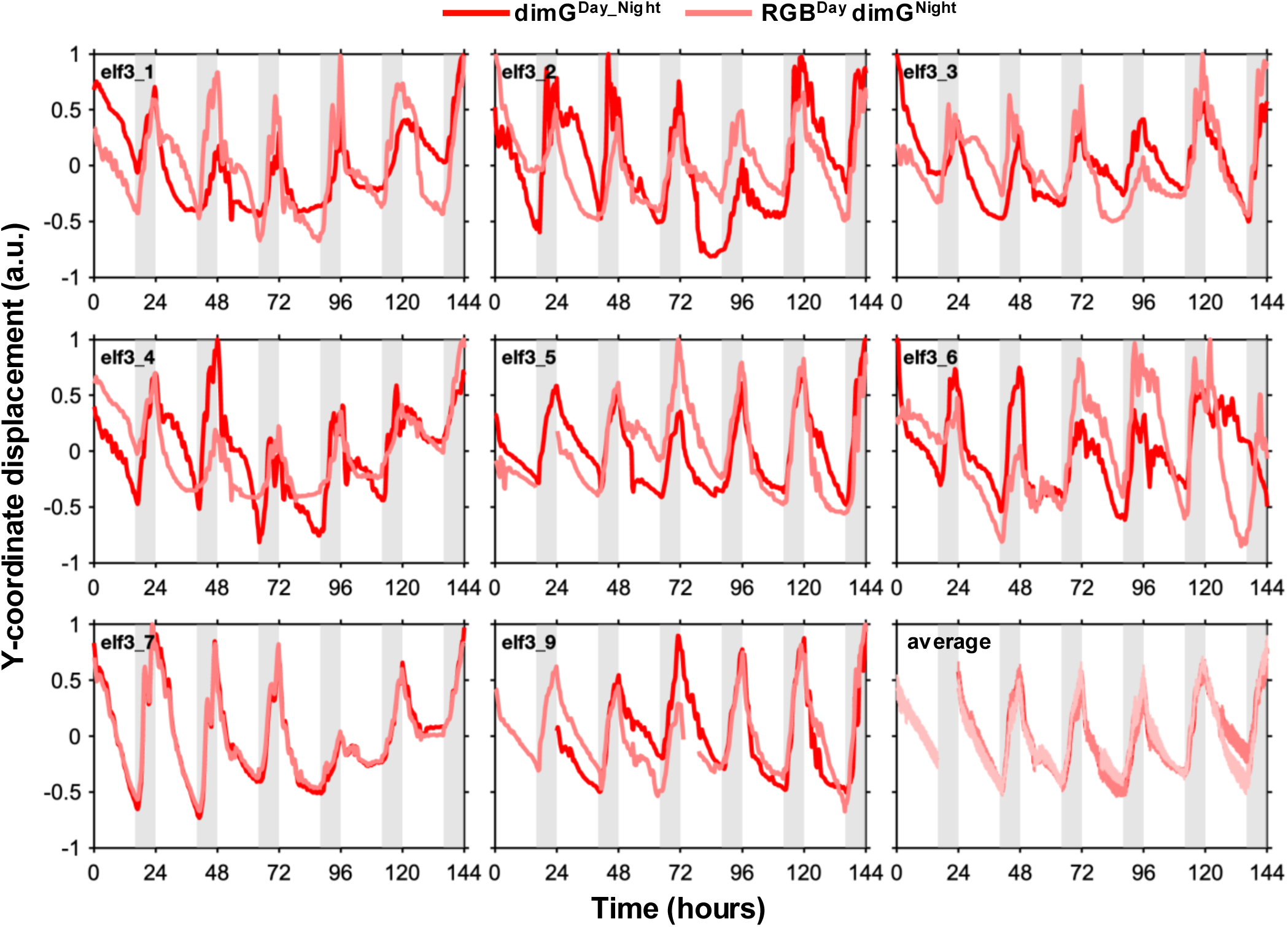
Diel oscillations of Y-coordinate displacement for individual plants using dimG^Day_Night^ and RGB^Day^_dimG^Night^ image sequences under LD. **(A)** Wild type, **(B)** *ztl*, **(C)** *lwd1 lwd2 and* **(D)** *elf3*.

**Figure S6.**
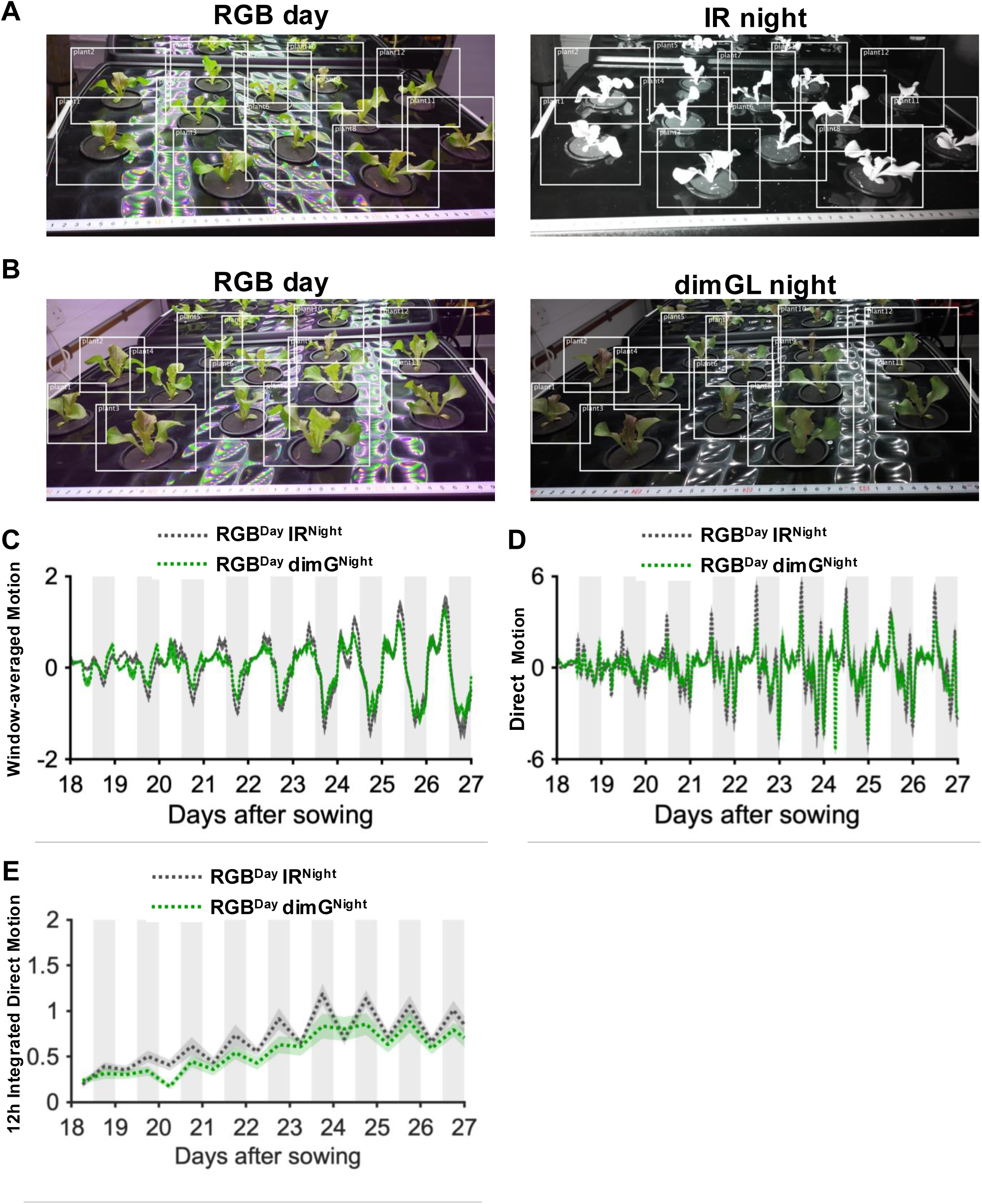
Comparison of IR vs dimG for night imaging, second replicate dataset. **(A-B)** Representative images for RGB^Day^ IR^Night^ (A) and RGB^Day^ dimG^Night^ (B) image sequences. **(C)** Mean Window-averaged Motion (dotted lines) ± SEM (shaded regions) for RGB^Day^ IR^Night^ (grey) and RGB^Day^ dimG^Night^ (green) conditions. **(D)** Mean Direct motion (dotted lines) ± SEM (shaded regions) for RGB^Day^ IR^Night^ (grey) and RGB^Day^ dimG^Night^ (green) conditions. **(E)** Mean 12-hour Integrated Motion (dotted lines) ± SEM (shaded regions) for RGB^Day^ IR^Night^ (grey) and RGB^Day^ dimG^Night^ (green) conditions. White and grey shaded in E-F panels indicate day and night, respectively.

**Figure S7.**
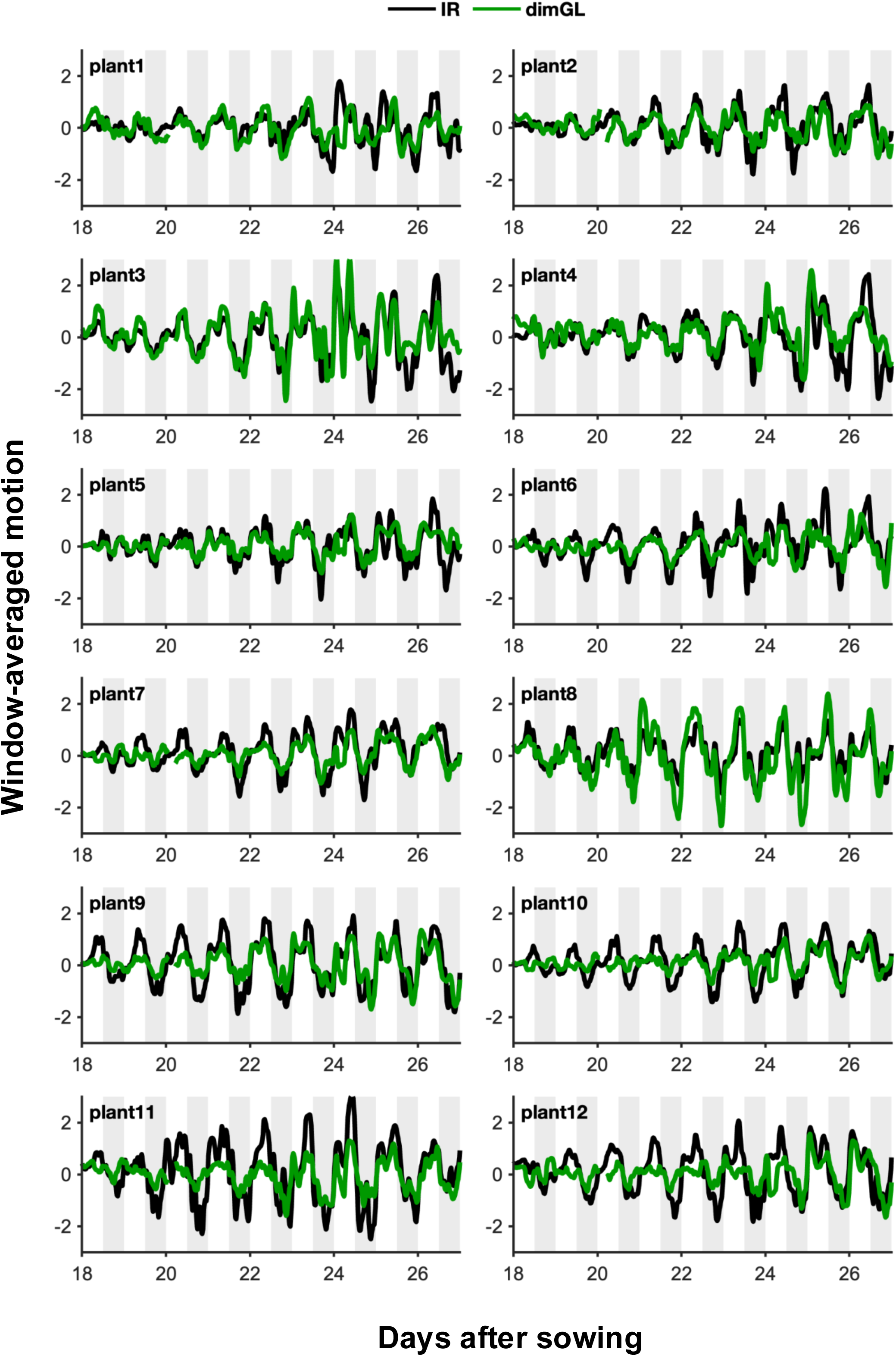
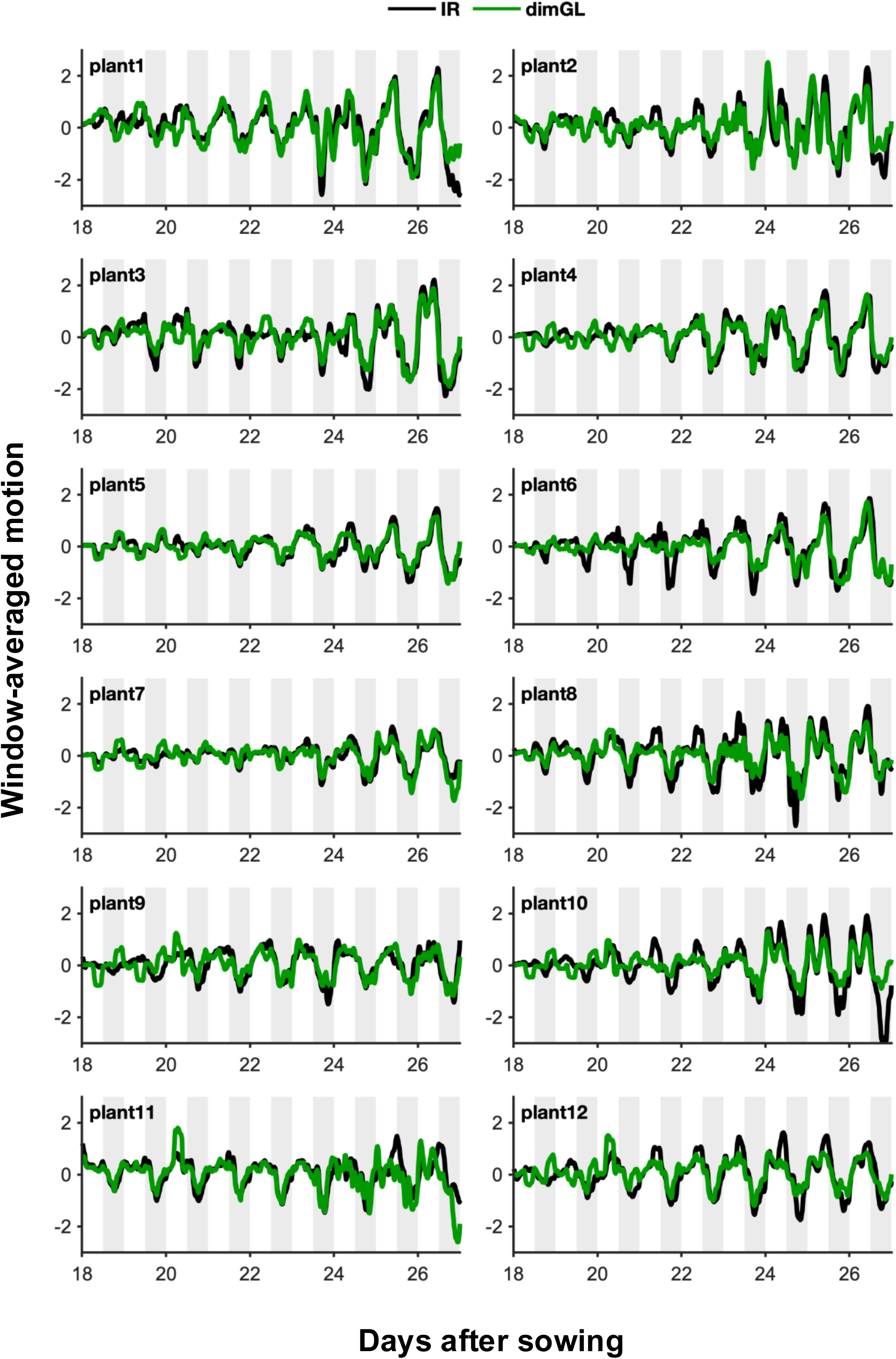
Window-averaged motion comparison of IR vs dimG for night imaging. Window-averaged motion for individual plants: (A) replicate 1 from Fig. 4D and (B) replicate 2 from Fig. S6C). IR night-imaging (black) and dimG nighttime-imaging (green). White and grey shaded areas indicate day and night, respectively.

**Figure S8.**
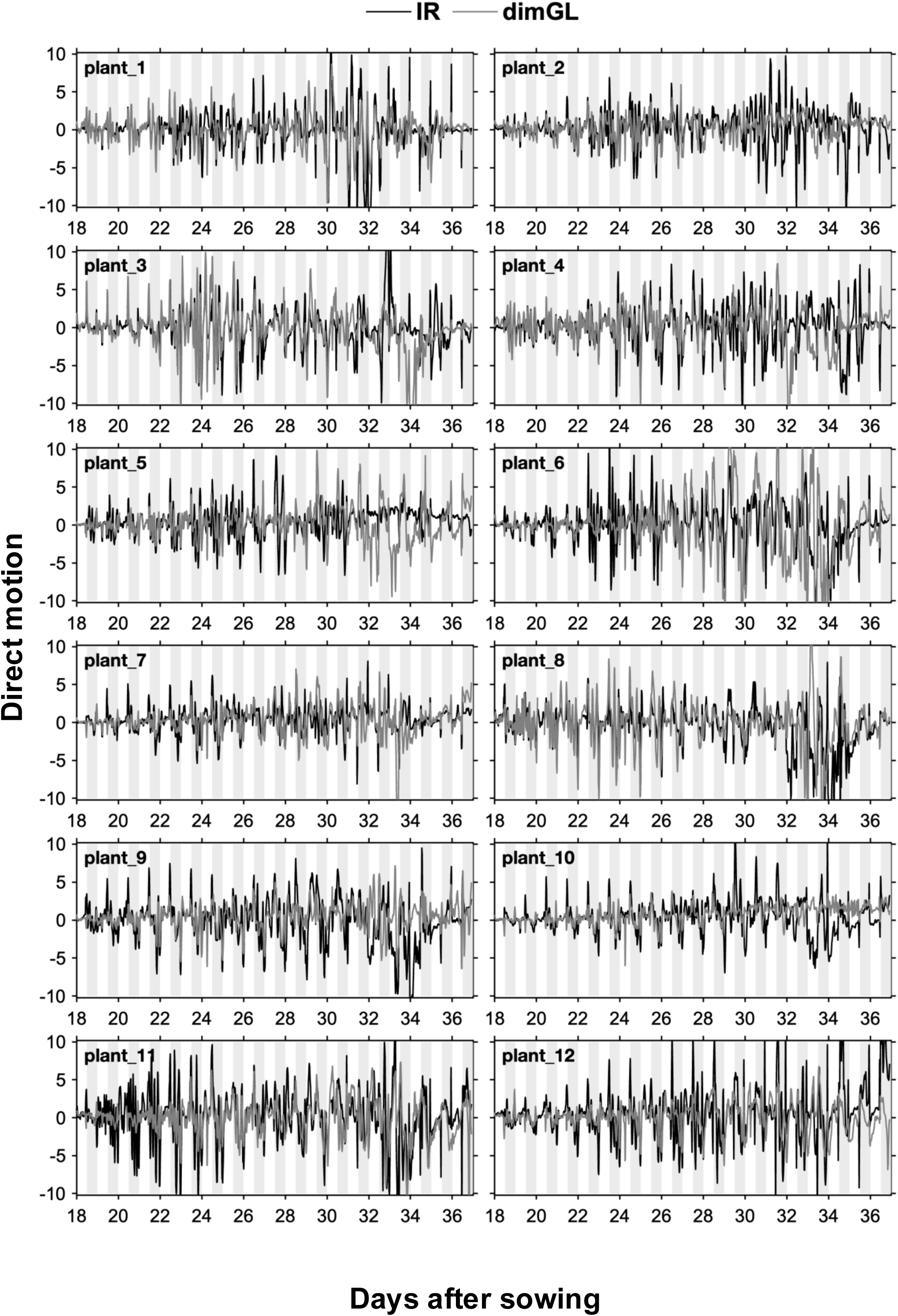
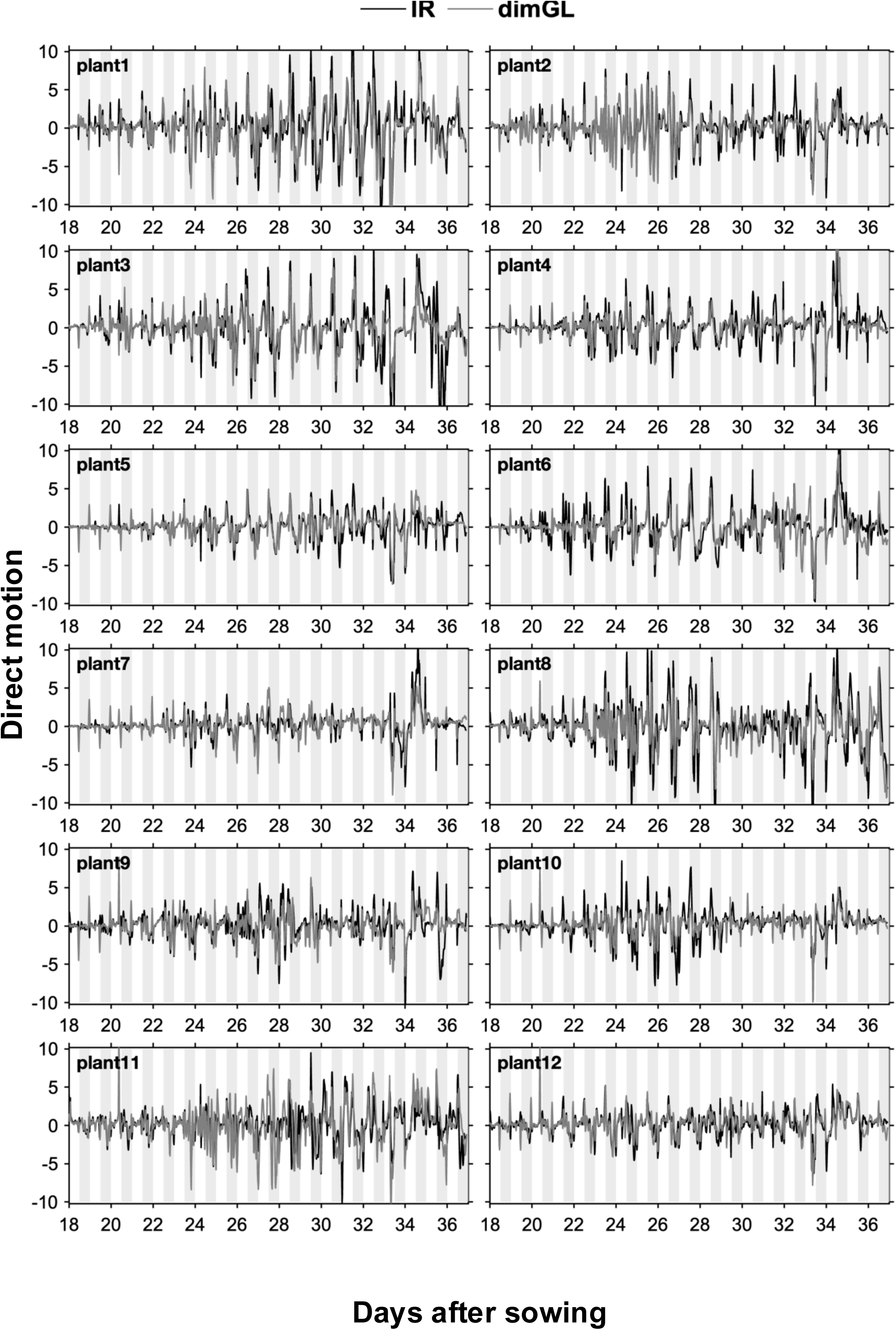
Direct motion comparison of IR vs dimG for night imaging. Direct Motion plots for individual plants: (A) replicate 1 from Fig.4E and (B) replicate 2 from Fig. S6D). IR night-imaging (black) and dimG night-imaging (grey). White and grey shaded areas indicate day and night, respectively.

**Figure S9.**
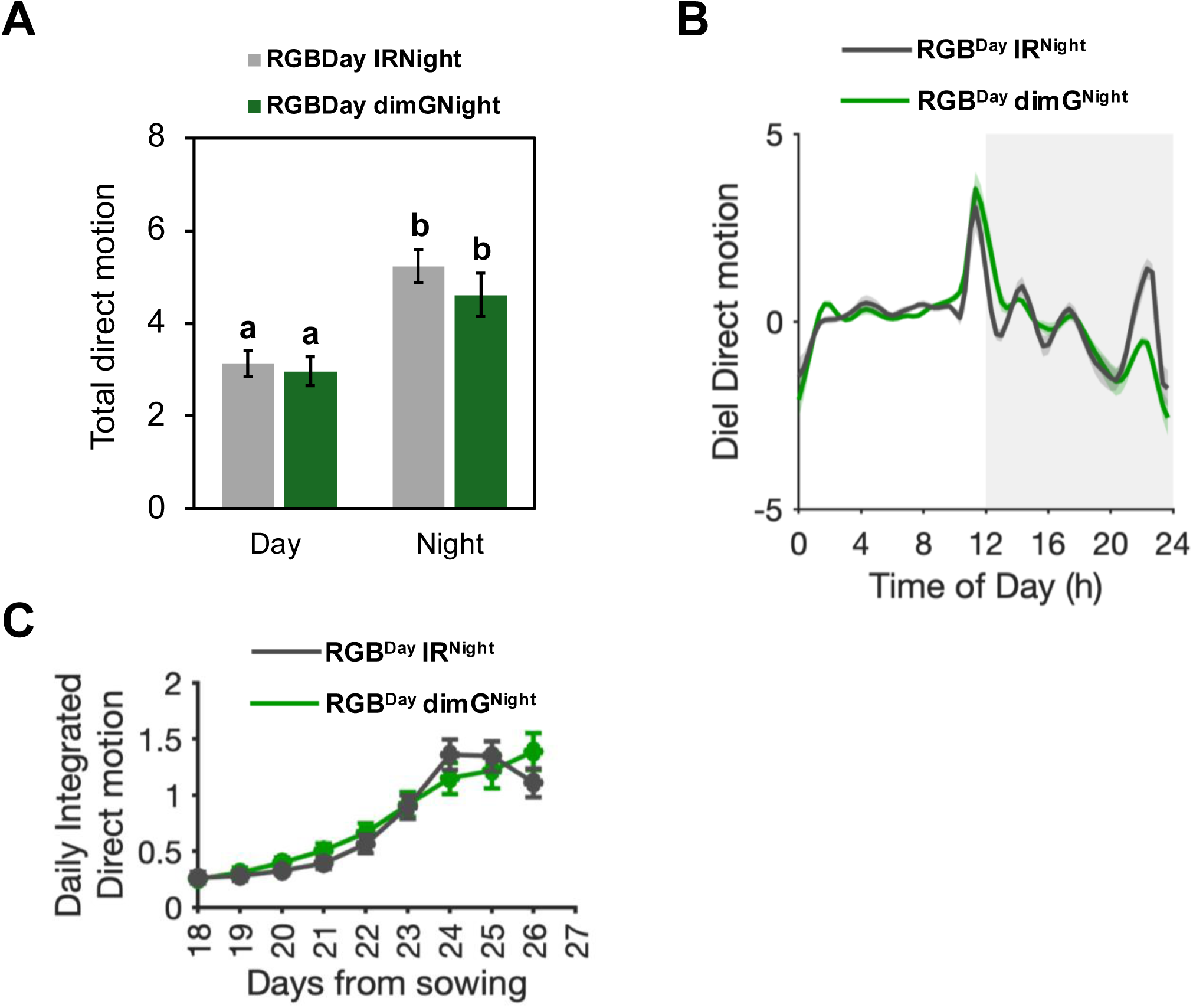
Higher leaf motion at night and increasing trend across the experiment. Extended analysis of data from Fig. 5B-C. **(A)** Total direct motion during day or night across the experiment (different letters, a and b, indicate groups with statistically similar means, Wilcoxon signed-rank test, V=78 n=12, *p* < 0.001). **(B)** Diel direct motion for days 18-27 (shading indicates SEM). **(C)** Daily integrated direct motion. For all panels (A–C), plants were grown under RGB^Day^ IR^Night^ (grey) or RGB^Day^ dimG^Night^ (green) conditions. Data represent means of 9–12 plants; error bars indicate SEM. Direct motion plots for individual plants associated are shown in Fig. S11A-B.

**Figure S10.**
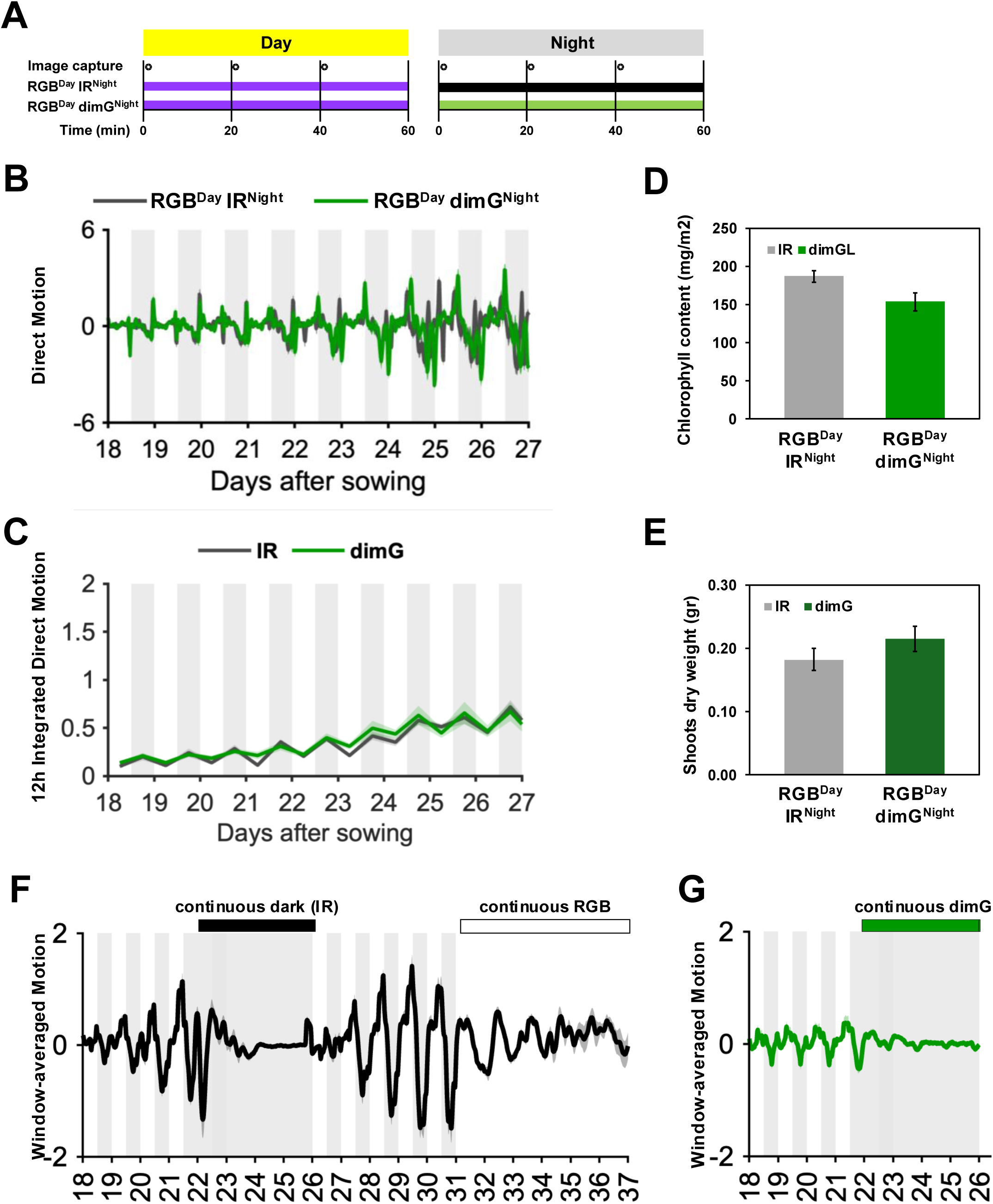
Nighttime exposure to continuous dimG does not influence leaf motion or overall plant growth in lettuce, second replicate dataset. **(A)** Imaging and lighting schedule. Plants were imaged every 20 minutes at the indicated times. During the day, full-spectrum RGB illumination was used (purple line); during the night, either infrared (IR; black line) or dim green (dimG; green line) illumination was applied. The photoperiod consisted of 12h day and 12h night. **(B)** Mean Direct Motion + SEM for lettuce plants (n=12) grown under RGB^Day^ IR^Night^ (grey) and RGB^Day^ dimG^Night^ (green) conditions. White and grey shading indicate day and night periods, respectively. **(C)** Mean 12-h Integrated Direct Motion (lines) ± SEM (shaded regions) for lettuce plants (n=12) grown under RGB^Day^ IR^Night^ (grey) and RGB^Day^ dimG^Night^ (green) conditions. White and grey shading indicate day and night, respectively. **(D-E)** Average chlorophyll content **(D)** and shoot dry weight **(E)** of lettuce plants grown under RGB^Day^ IR^Night^ (grey) and RGB^Day^ dimG^Night^ (green) conditions, measured at day 28. Error bars indicate SEM, n=12. **(F)** Continuous light, but not continuous darkness, sustains leaf movement oscillations in lettuce. Mean Direct Motion + SEM for lettuce plants (n=12) grown under RGB^Day^ IR^Night^ (days 18-21 and 26-29), continuous darkness (IR; days 22-25) and continuous light (RGB; days 30-37). White and grey shading indicate light (RGB) and dark (IR) conditions, respectively. **(G)** Continuous dimG does not sustain leaf movement oscillations. Mean Direct Motion + SEM for lettuce plants grown under RGB^Day^ dimG^Night^ (days 18-21) and continuous dimG (days 22-25). White and grey shading indicate light (RGB) and dark (dimG), respectively. Direct motion plots for individual plants associated with panels C–D are shown in Fig. S11C–D, and those associated with panels F–G are shown in Fig. S12B–C.

**Figure S11.**
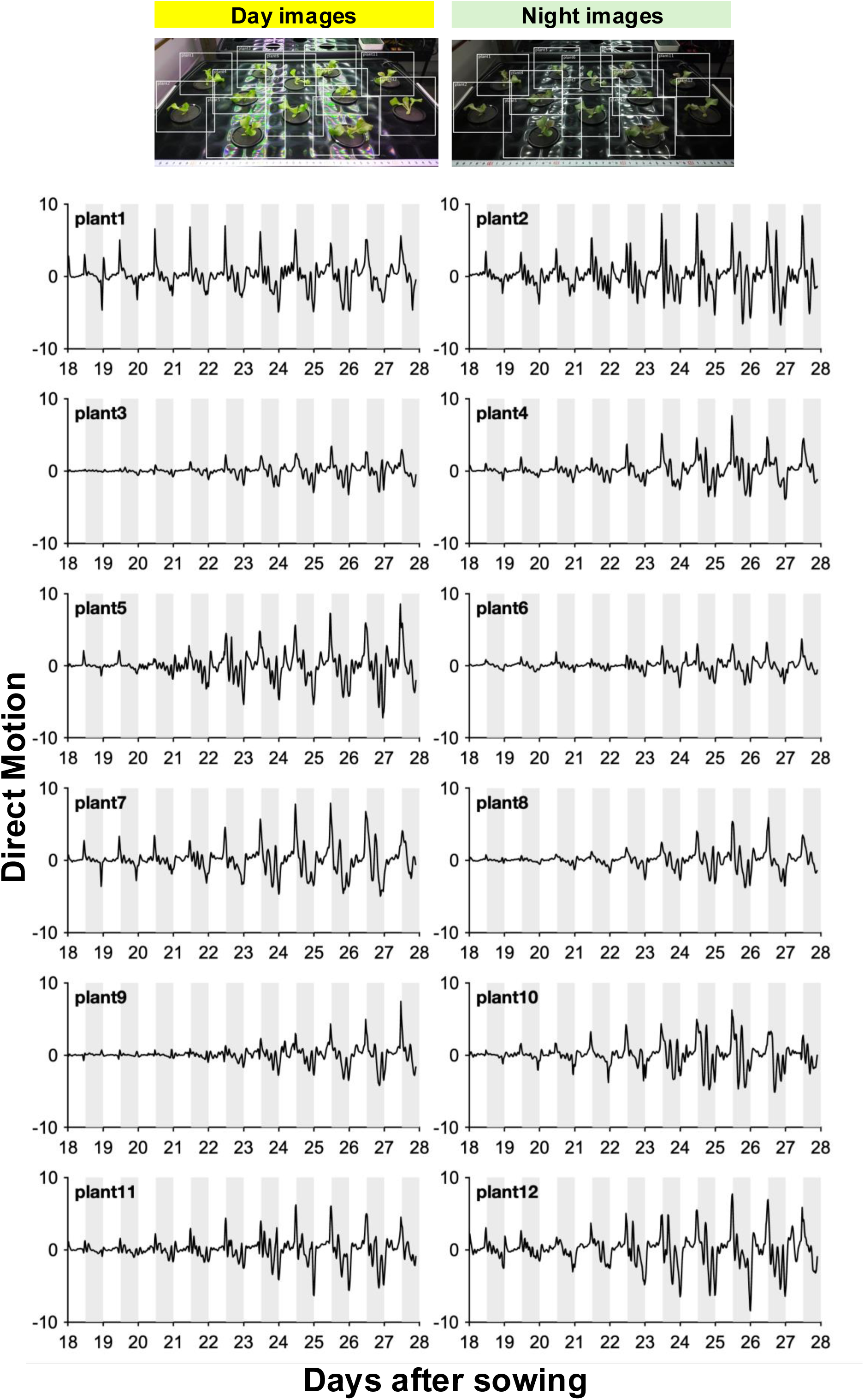
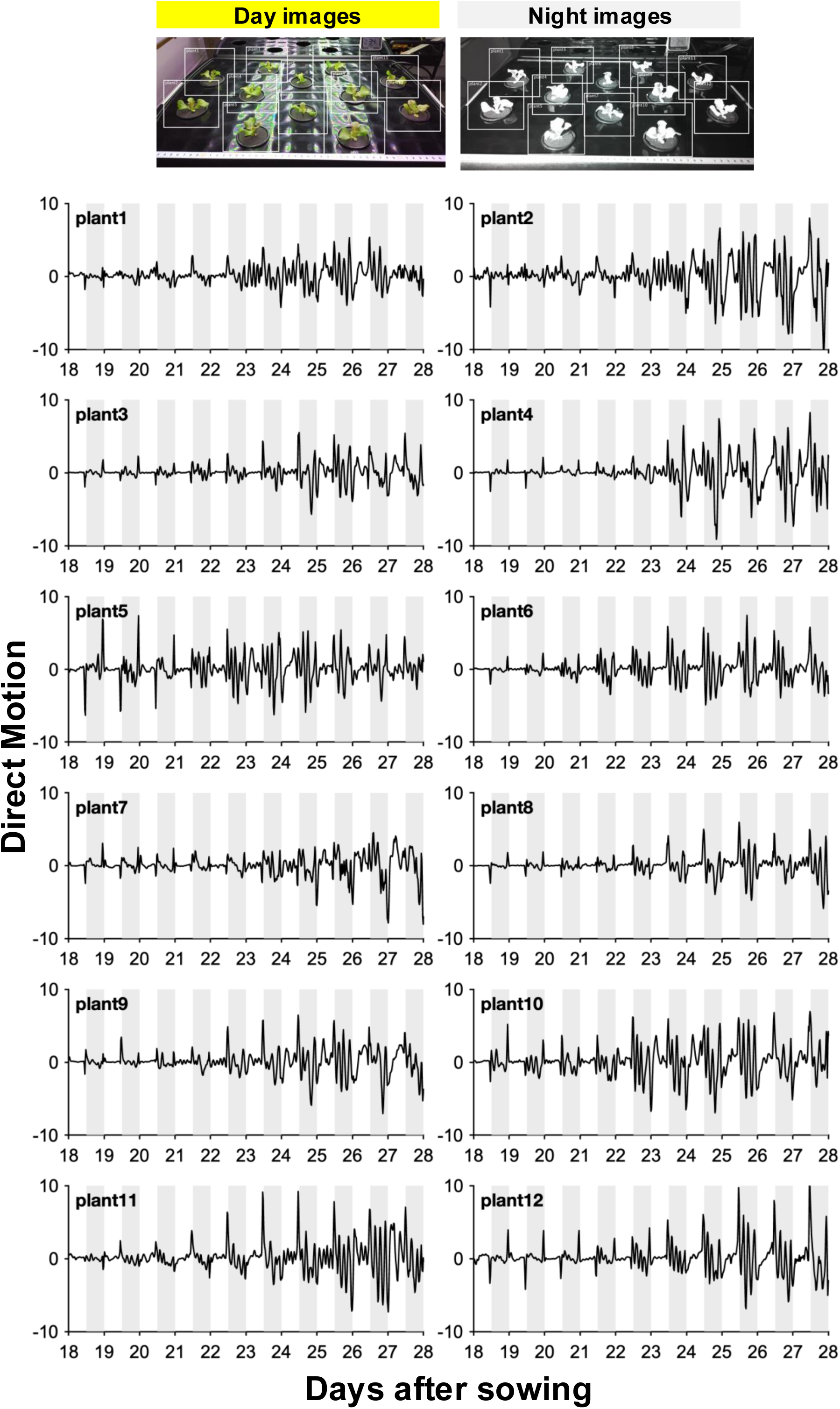
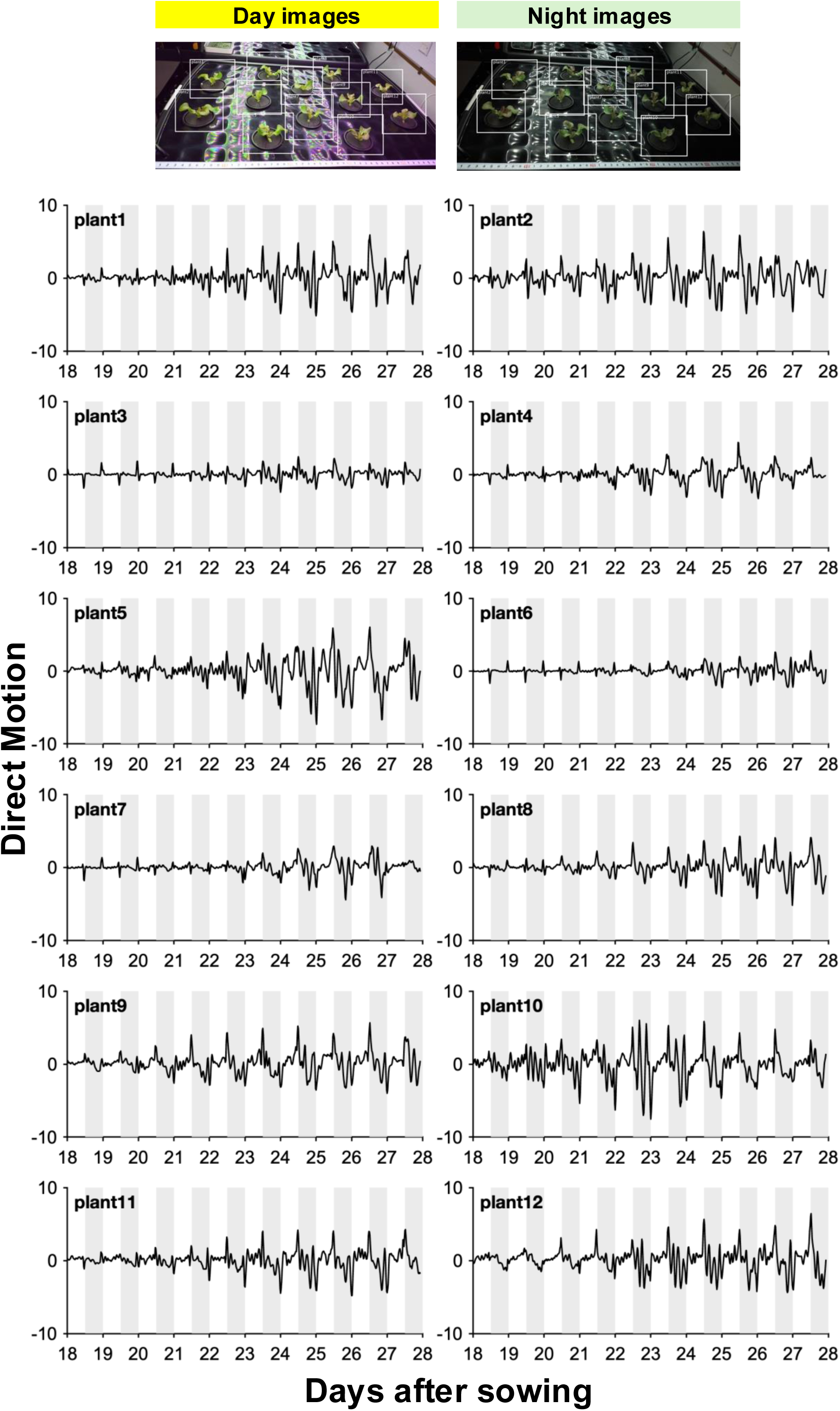
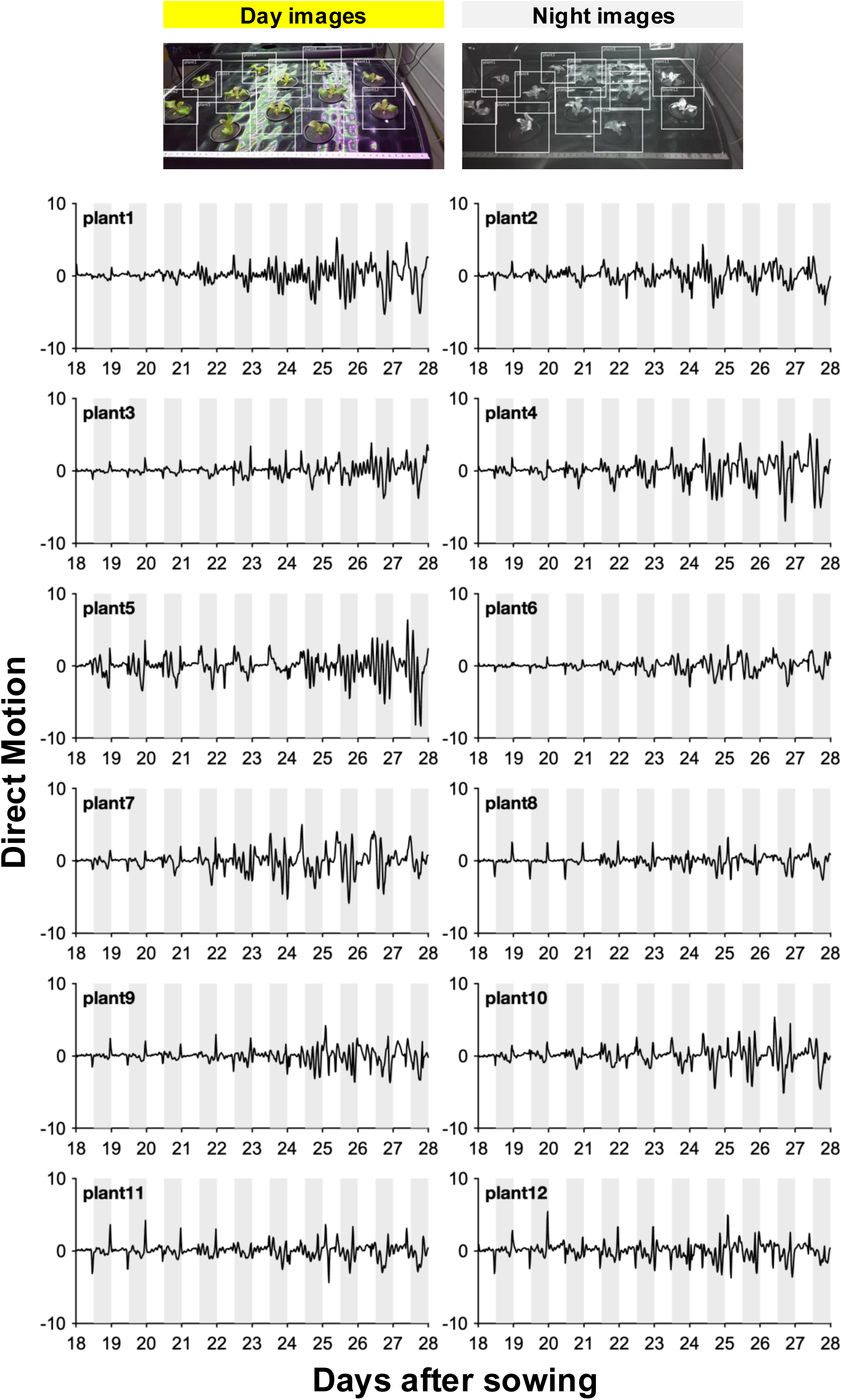
Direct motion of plants grown under continuous dimG or under darkness (IR) during the night. Representative images depicting ROIs used for TRiP motion estimation and Direct motion plots for individual plants: Replicate 1 from Fig. 5 B-C **(A)** continuous dimG at night and **(B)** continuous darkness (IR). Replicate 2 from Fig. S10 B-C **(C)** continuous dimG at night and **(D)** continuous darkness (IR). White and grey shading indicate light (RGB) and dark (dimG), respectively.

**Figure S12.**
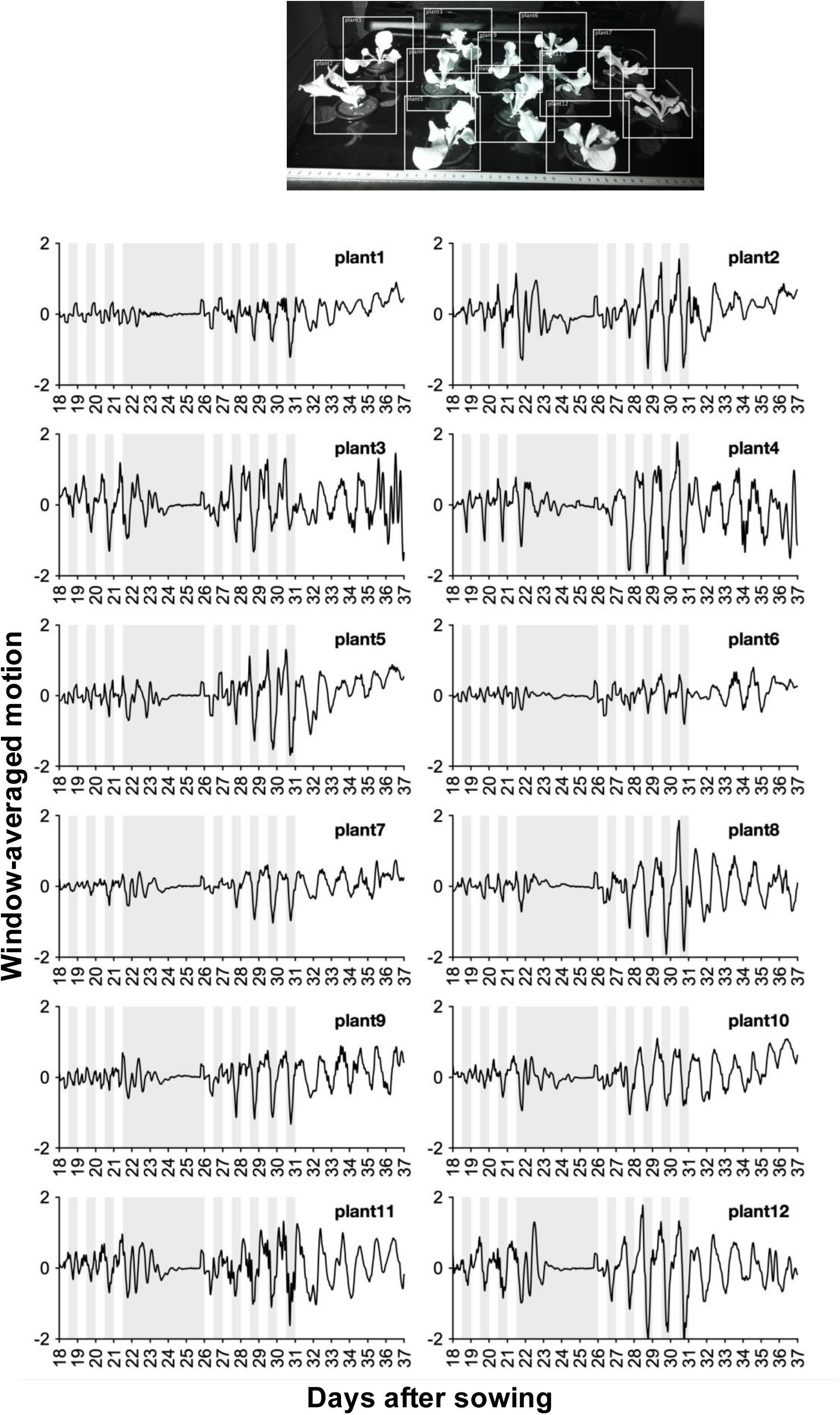
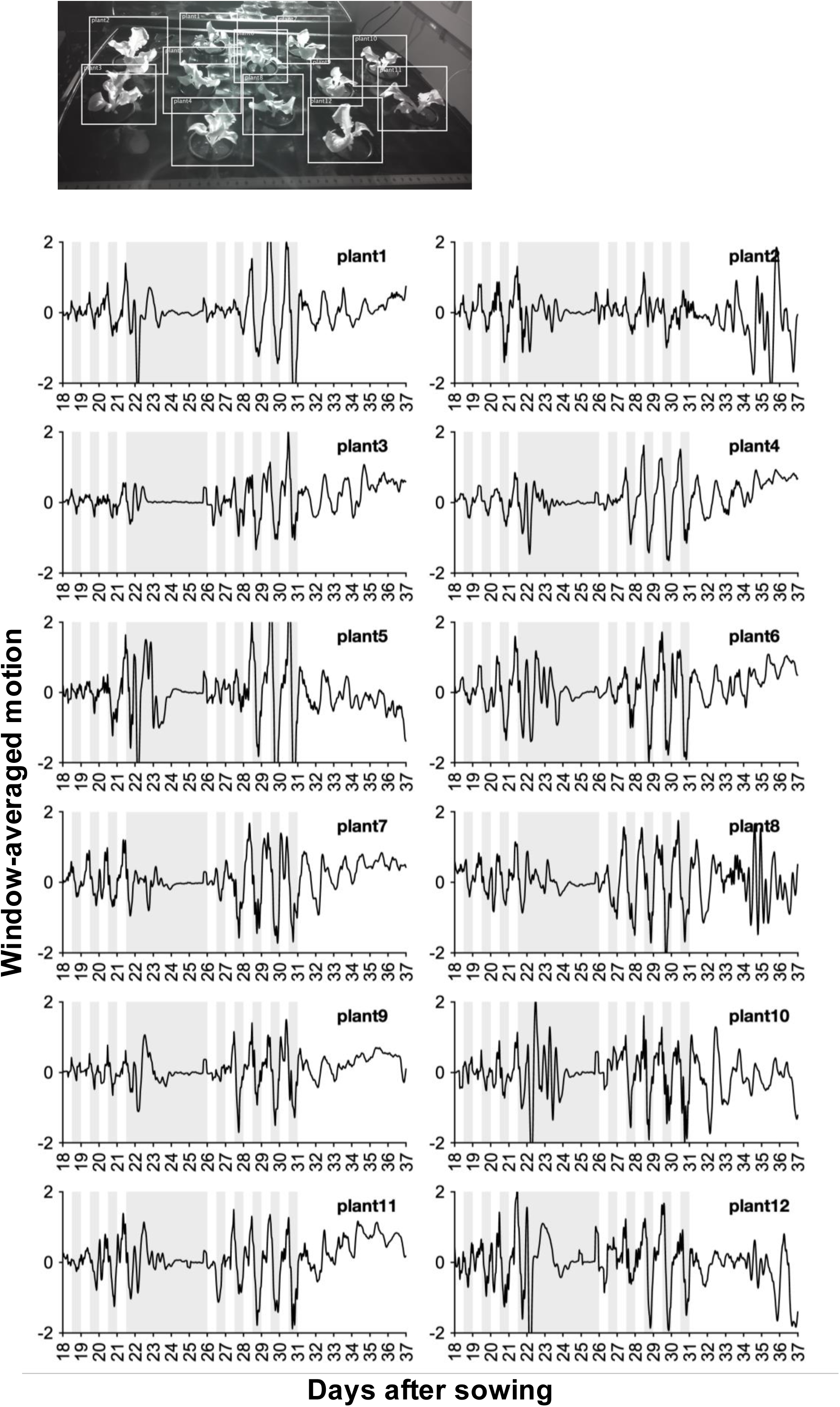
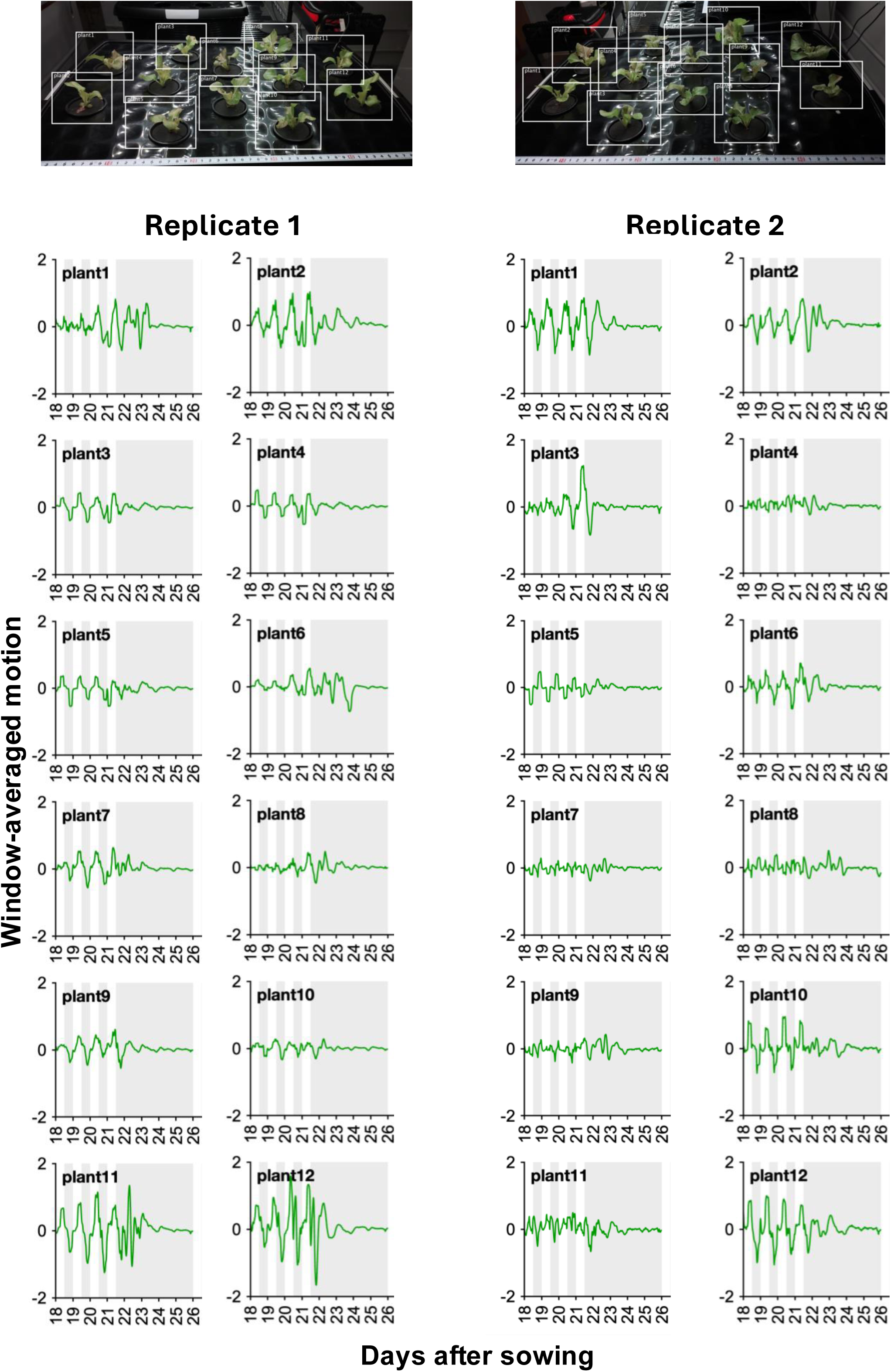
DD and dimG do not sustain leaf movement under free-running conditions. Representative images depicting ROIs used for TRiP motion estimation, and Window-averaged motion for individual plants: **(A)** DD and LL experiment (data from Fig. 5F), **(B)** DD and LL experiment (replicate 2 Fig. S10F, **(C)** continuous dimG: replicate 1 from Fig. 5G (left) and right data from replicate 2 Fig. S10G (right). White and grey shading indicate light (RGB) and dark (dimG), respectively.

